# Paired Deep Mutational Scanning of Lck Activity and Inhibitor Sensitivity Enables the Engineering of Pharmacological Control of T Cell Receptor Signaling

**DOI:** 10.64898/2026.11.13.766442

**Authors:** Zachary E. Potter, Amber N. Graves, Jordan L. Brown, Cindy T. Wei, Jorge Cardenas, Melissa A. Chiasson, Jessica J. Simon, Nicholas A. Popp, Daniel S. Brush, Doria E. Unrau, Douglas M. Fowler, Dustin J. Maly

**Author notes:** Correspondence (D.J.M.).

## Abstract

Controlling signaling enzymes with small molecules is a central goal in chemical biology and is of great utility for engineering cell-derived immunotherapies. Yet strategies for conferring selective small-molecule sensitivity without disrupting a protein’s native function remain limited. Here we present a generalizable approach to engineering small molecule-controlled multidomain protein kinases. Using deep mutational scanning of full-length Lck, a tyrosine kinase essential for T cell receptor (TCR) signaling, we measured the effect of nearly all single amino acid substitutions in the kinase domain on phosphotransferase activity. The resulting activity landscape defined the domain’s tolerance to substitution, mapped the regulatory interfaces that tune phosphotransferase activity, and, by treating bulky substitutions as ligand proxies, nominated cryptic pockets where covalent liganding is predicted to allosterically inhibit Lck. In parallel screens, we measured the sensitization of each variant to five clinically approved, reversible drugs: dabrafenib, erlotinib, gefitinib, imatinib, and nilotinib. Together, these landscapes identified substitutions that preserved or enhanced activity while sensitizing Lck to inhibition. Sensitizing substitutions were distributed throughout the kinase domain, including positions more than 30 Å from the ATP-binding site, implicating long-range allosteric networks linking distal perturbations to ATP-binding site conformation and inhibitor potency. Integrating the activity and sensitization landscapes yielded a nilotinib-sensitized variant, I312S, that supports TCR signaling and, upon drug treatment, selectively suppresses proximal signaling and IL-2 production in Jurkat cells even in the presence of endogenous Lck.

## Introduction

Intracellular multidomain signaling enzymes govern cell fate decisions and are attractive targets for both small-molecule drug discovery and cell-based therapeutic engineering. Small molecules remain the predominant modality for controlling protein function in biology and medicine, with the vast majority of FDA-approved drugs acting on protein targets (Santos, et al., 2017; Southey and Brunavs, 2023), yet our ability to engineer proteins that retain native function while acquiring selective drug sensitivity is still limited. Combining function-preserving mutations with substitutions that confer sensitivity to FDA-approved small molecules offers a powerful route for the selective control over engineered proteins and their cellular outputs. Identifying permissive sites in large, conformationally dynamic enzymes remains difficult, however; zero-shot prediction algorithms lack sufficient accuracy, while more rigorous force-field-based approaches are computationally intractable at scale (Liu, et al., 2025). Large-scale mutational screens offer a reliable experimental alternative for mapping the sequence-function landscapes that guide rational protein engineering.

The Src-family kinase Lck is particularly well suited to such efforts. Lck plays a central, non-redundant role in T cell receptor (TCR) signaling (Straus and Weiss, 1992), where precise tuning of its activity governs signal amplification and tolerance to self-pMHC ligands (Horkova, et al., 2020). Its SH2 and SH3 domains, together with its kinase domain (KD), establish a conformational equilibrium between active and autoinhibited states (**Fig. S1A**), and this multidomain architecture offers multiple handles for tuning phosphotransferase activity and intermolecular scaffolding through sequence variation. Consistent with this, altering Lck activity or expression level profoundly affects T cell fate and function (Moline, et al., 1992; Abraham, et al., 1991; Levin, et al., 1993). Lck is thus a compelling target for both pharmacological control and rational engineering of T cell signaling (Sun, et al., 2020; Weber, et al., 2021).

Here, we performed a deep mutational scan of Lck’s KD in the context of the full-length enzyme, quantifying the functional consequences of nearly all single amino acid substitutions. This approach revealed a broad landscape of substitutions that modulated phosphotransferase activity, including an enrichment of deactivating substitutions proximal to certain endogenous lysine amino acids, pointing to cryptic ligandable pockets within Lck’s KD. These observations address a persistent challenge in covalent ligand discovery: determining whether liganding a given nucleophilic position will perturb the target protein’s function. Focusing on cysteine and lysine substitutions that retained kinase activity, we identified a cysteine variant that confers sensitivity to the covalent, FDA-approved EGFR inhibitor neratinib. By repurposing our activity screen into an inhibitor sensitization platform, we found additional substitutions that render Lck sensitive to the reversible, ATP-competitive inhibitors dabrafenib, erlotinib, gefitinib, imatinib, and nilotinib. Integrating these datasets yielded a nilotinib-sensitized Lck variant that functions as a small molecule-controlled “off switch” for TCR signaling in Jurkat cells. Together, these results demonstrate that pairing deep mutational scanning of phosphotransferase activity with measurements of drug sensitization provides a generalizable route to engineering pharmacologically controllable signaling enzymes, with principles likely extending beyond Lck to other kinases and multidomain enzymes that function through complex regulatory networks.

## Results

### Activity measurements for 4,892 Lck variants

To assess the activity of Lck variants, we turned to a yeast-based growth assay that exploits the inverse correlation between Src-family kinase phosphotransferase activity and *S. cerevisiae* growth (**Fig. 1A**). Yeast provides a eukaryotic cellular environment devoid of extrinsic regulatory factors in which to profile tyrosine kinase activity (Ahler, et al., 2019; Brugge, et al., 1987). As observed for Src, yeast expressing a kinase-dead Lck variant grew faster than yeast expressing WT Lck, whereas yeast expressing activating Lck variants grew most slowly (**Fig. 1A**). To systematically define how sequence variation within Lck’s KD affects activity, we performed a deep mutational scan of the KD (positions 245-498) in the context of full-length Lck, leaving all regulatory domains intact. Because Lck’s SH2 and SH3 domains mediate interactions essential to TCR activation (Lo, et al., 2018), we reasoned that restricting variation to the KD would best allow us to tune enzymatic activity, signaling behavior, and drug sensitivity while preserving the scaffolding required for native TCR signaling, and for the eventual deployment of engineered Lck variants in T cell-based systems.

**Figure 1.**
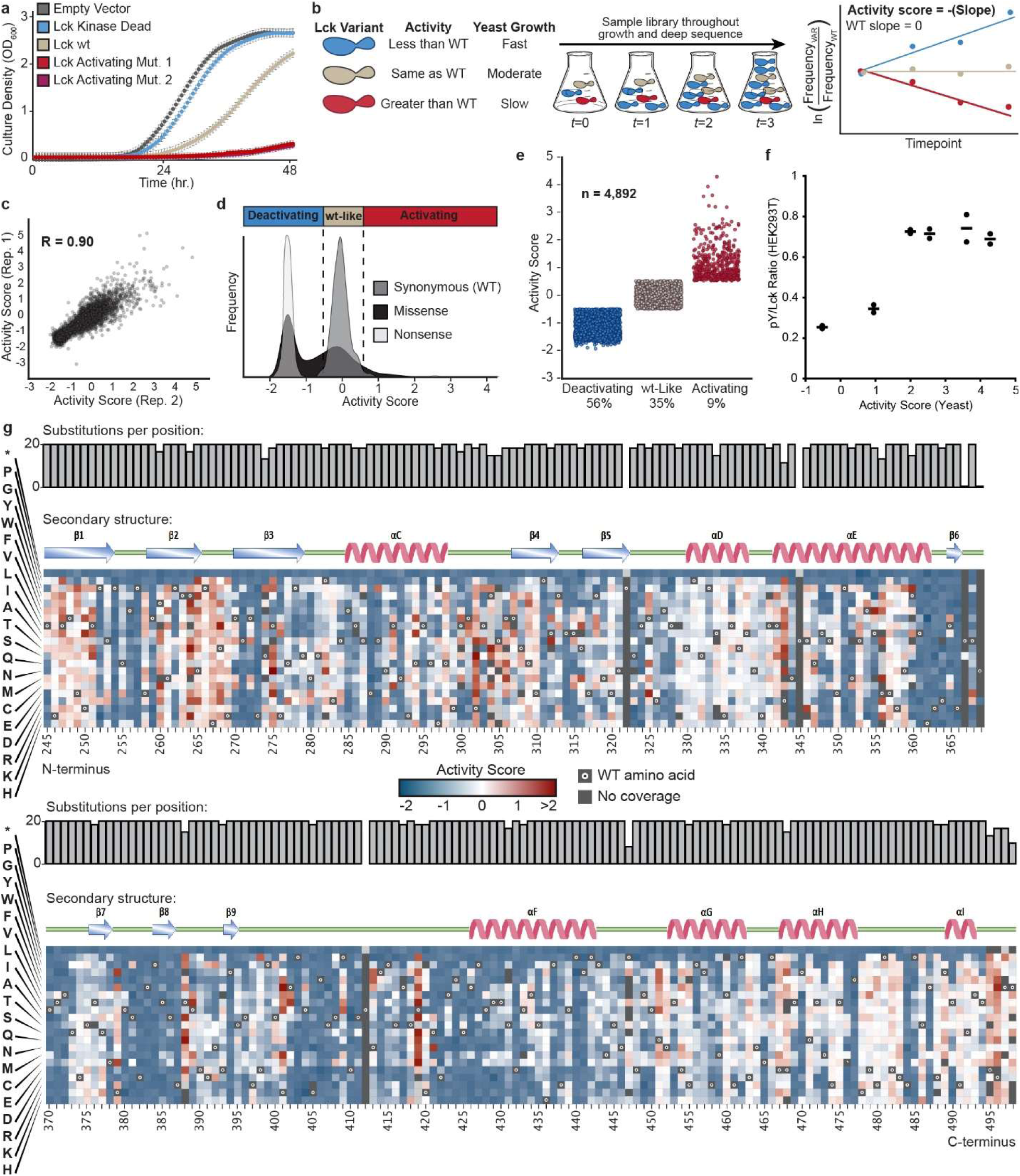
Kinase Activity Measurements of 4,892 Single Amino Acid Variants of Lck. (A) Individually assessed growth curves for yeast expressing Lck WT, K273R (kinase dead), one of two activating variants, or a control vector. Datapoints represent average ± s.e.m. of three biological replicates. (B) Schematic for yeast growth-based deep mutational scan of Lck’s KD. (C) Scatterplot showing correlation between activity scores calculated from two independent biological replicates of the Lck variant library. (D) Histogram of activity scores for synonymous (WT), non-synonymous, or missense variants. The dashed lines indicate two standard deviations from the average of the synonymous (WT) distribution, which we used to define Activating (right or rightmost dashed line) and Deactivating (left of leftmost dashed line) variants. (E) Activity scores for variants classified as Deactivating (n=2,714), wt-Like (n=1,720), or Activating (n=458). (F) Scatter plot of activity scores calculated in yeast as they correspond to activation loop phosphotyrosine (pY394) levels in HEK293T cells. Datapoints represent two separate biological replicates, bars represent mean. (G) Sequence-activity map for all positions in Lck’s KD. Each column represents a position in Lck’s KD (starting at position 245 and ending at position 498) and each row represents an amino acid substitution (* = nonsense mutation). Black dots represent the wild-type amino acid and grey tiles indicate missing data. Bar graphs (Substitutions per position) represent mutational completeness at each amino acid position. Secondary structure and functional motif annotations were obtained from the ProKinO database.

We had a DNA-barcoded plasmid library of full-length Lck commercially synthesized in which every KD position was substituted with all other amino acids and with a stop (nonsense) codon, with Lck expression under the control of a galactose-inducible promoter. PacBio long-read sequencing of the plasmid library was used to associate individual DNA barcodes with individual Lck variants, yielding a library of 4,959 variants (97.6% mutational completeness), with each variant encoded by an average of 29 barcodes (**Fig. S1B**). We transformed the library into yeast and collected samples before Lck induction and at successive timepoints during outgrowth (**Fig. 1B**). Plasmids were extracted at each timepoint, and next-generation sequencing (NGS) was used to quantify the relative frequency of each barcode and thereby each variant. Using a weighted least squares scoring method from the Enrich2 software package (Rubin, et al., 2017), we calculated activity scores for 4,892 of the 4,959 variants in our plasmid library (4,646 missense and 246 nonsense) by averaging two highly correlated independent biological replicates (r = 0.90; **Fig. 1C**). Scores were normalized to the distribution of synonymous WT variants, which served as an internal reference for WT kinase activity. This distribution was tightly clustered and well separated from deactivating nonsense variants, confirming that variants with WT-like activity could be resolved from those with lower activity. We defined a variant as WT-like if its activity score fell within two standard deviations of the synonymous WT mean, and as activating or deactivating if it fell more than two standard deviations above or below that mean, respectively. By this definition, 35% of variants had WT-like activity, 9% were activating, and 56% were deactivating (**Fig. 1D** and **1E**).

To validate our activity screen, we measured growth curves of yeast individually expressing Lck variants and found that these individual growth curves closely mirrored the activities measured in our deep mutational scan: variants with high activity scores displayed slowed growth relative to WT, and variants with low activity scores grew faster than WT (**Fig. S1C**). We also measured activation loop phosphorylation levels for six Lck variants transiently expressed in HEK293T cells and found that these levels corresponded with calculated activity scores (**Fig. 1F**). Representation of our activity data as a sequence-activity map recapitulated known features of kinase structure-function relationships (**Fig. 1G**). For example, nearly all substitutions at positions K273, E288, H362, R363, D364, and D382, which make up the catalytic machinery of tyrosine kinases (Knighton, et al., 1991), were classified as deactivating. Additionally, all substitutions at the activation loop tyrosine Y394 were deactivating, consistent with its autophosphorylation being required for full catalytic activity. We identified 33 positions at which every substitution was deactivating, indicating that the identity of the WT amino acid at these positions is essential to Lck’s phosphotransferase activity. These positions are generally clustered around Lck’s ATP-binding site and form a network connecting the N-and C-lobes of the KD (**Fig. S1D**). We next examined the catalytic and regulatory spine positions, which together constitute the hydrophobic spine whose assembly is required for protein kinase catalytic activity (Kornev, et al., 2006; Kornev, et al., 2008). Consistent with their importance for maintaining Lck’s catalytic activity, more than 90% of substitutions were deactivating at 15 of the 20 spine positions.

### The KD of Lck is tolerant of amino acid substitution

Although 33 positions in the KD were completely intolerant of substitution, 45% of all missense variants retained WT-like or higher activity, indicating that the KD tolerates extensive variation while preserving catalytic function. To obtain a position-resolved map of mutational tolerance, we projected the number of WT-like and activating substitutions at each position onto the structure of Lck’s KD (**Fig. 2A, S2A**). We found that 86% of all positions in the KD had at least one WT-like or activating missense variant, and 63% tolerated five or more such substitutions (**Fig. 2A**). As expected, positions lining the ATP-binding site were the most constrained: only six of the thirteen positions (46%) within 5 Å of the site had five or more WT-like or activating substitutions, compared with 64% for more distal positions (>5 Å). Nevertheless, even 8 of the 13 positions (62%) lining the ATP-binding site harbored at least three WT-like or activating substitutions.

**Figure 2.**
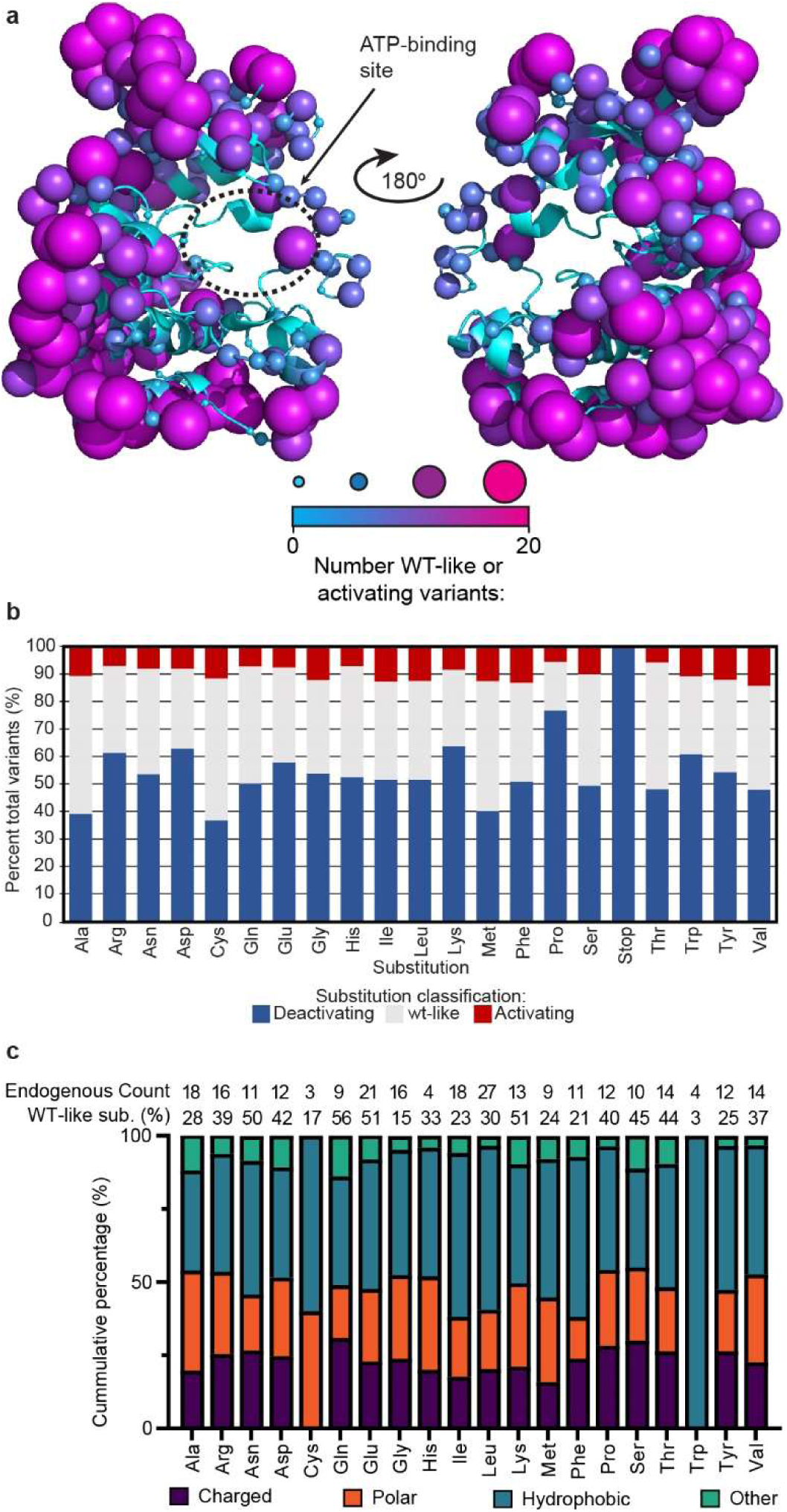
The KD of Lck is highly tolerant of substitution. (A) The number of WT-like and activating substitutions at each-position projected onto an AlphaFold model of Lck’s KD (AlphaFold: AF-P06239-F1). Sphere size and color represent the number of WT-like and activating amino acid substitutions observed at each position. (B) Stacked bar graph representing the percentage of each amino acid substitution’s activity score classification. (C) The number of occurrences of each amino acid type in Lck’s KD (top row) and the percentage of substitutions that are WT-like for each amino acid type (bottom row). Stacked bar graph represents the proportion of each amino acid substitution that is WT-like for an amino acid type (charged: Arg, Asp, Glu, His, Lys; polar: Asn, Gln, Ser, Thr; hydrophobic: Ala, Ile, Leu, Met, Phe, Trp; or other: Gly, Pro) in Lck’s KD (x-axis).

We next asked which substitutions most often preserve WT-like activity (**Fig. 2B**) and which wild-type amino acid types are most tolerant of substitution (**Fig. 2C**). Ala substitutions were among the best tolerated, with 50% retaining WT-like activity, but Cys substitutions were slightly better tolerated (52%), and Met substitutions were close behind (47%). As expected, Pro was the least tolerated substitution, though 18% of Pro substitutions retained WT-like activity. The charged substitutions Lys, Arg, and Asp were the next least tolerated. Grouped instead by WT amino acid identity, positions occupied by Glu or Lys were the most tolerant of substitution, with 51% of variants at these positions retaining WT-like activity, consistent with a broader trend in which polar positions show the highest tolerance, likely reflecting their tendency to be solvent-exposed rather than buried. Conversely, the rarest WT amino acids in the KD (Trp, 4 occurrences; Cys, 3; His, 4) were the least tolerant: Trp (4 positions, 3% of substitutions WT-like), Cys (3, 17%), and His (4, 33%). Notably, Cys is readily accommodated as a substitution yet is itself poorly replaced, as expected if the KD’s three endogenous Cys positions serve specific structural or functional roles. Among amino acids occurring at more than five positions, Met, Ile, and Phe were the least tolerant of substitution, consistent with a general trend in which buried hydrophobic positions show the lowest tolerance, likely because few substitutions reproduce their favorable packing interactions. Gly was similarly intolerant, though most likely because no other amino acid can recapitulate the conformational flexibility and minimal steric footprint that Gly often provides. Together, these data map mutational tolerance across Lck’s KD, distinguishing the small set of positions that cannot be varied from the majority that accommodate substitution while preserving phosphotransferase activity.

### Substitutions at regulatory interfaces tune the allosteric modulation of Lck’s phosphotransferase activity

We observed that 9% of all variants within Lck’s KD had increased phosphotransferase activity relative to WT Lck (**Fig. 1E**). Such activating variants were previously observed to occur in the closely related tyrosine kinase Src at domain-domain interfaces within the KD that mediate intramolecular autoinhibition (Ahler, et al., 2019; Beltran, et al., 2026). To determine whether the same holds for Lck, and thus whether varying interface positions offers a way to tune phosphotransferase activity while leaving the scaffolding functions of the regulatory domains intact, we generated a structure-activity map by projecting position-averaged activity scores onto an AlphaFold model of Lck’s SH3, SH2, and KD (positions 73-509; **Fig. 3A**). This analysis revealed spatially clustered regions in which multiple substitutions increased phosphotransferase activity, reminiscent of patterns observed in Src (Ahler, et al., 2019; Beltran, et al., 2026), suggesting that these regions likewise correspond to interfaces mediating the domain-domain interactions that modulate Lck autoinhibition. To define these regions systematically, we selected KD positions harboring five or more activating substitutions (34 positions) and performed hierarchical clustering on their α-carbon coordinates in the AlphaFold model of Lck (**Fig. 3B**). This yielded ten discrete clusters, seven of which mapped onto previously defined regulatory regions in Src, including the SH2:KD interface, the SH3/linker:KD interface, the αF pocket, and an N-terminal lobe cluster associated with kinase hyperactivation and drug resistance (Ahler, et al., 2019; Chakraborty, et al., 2024). Notably, the SH3/linker:KD interface (clusters 3 and 4) and the SH2:KD interface (clusters 8 and 10) showed the greatest divergence from Src, with Lck exhibiting both a higher density and a greater magnitude of activating substitutions at these surfaces despite equivalent coverage in each DMS dataset. This suggests that Lck’s kinase activity is particularly sensitive to perturbation of allosteric coupling between the KD and the SH3-SH2 module (**Fig. 3A-3C**).

**Figure 3.**
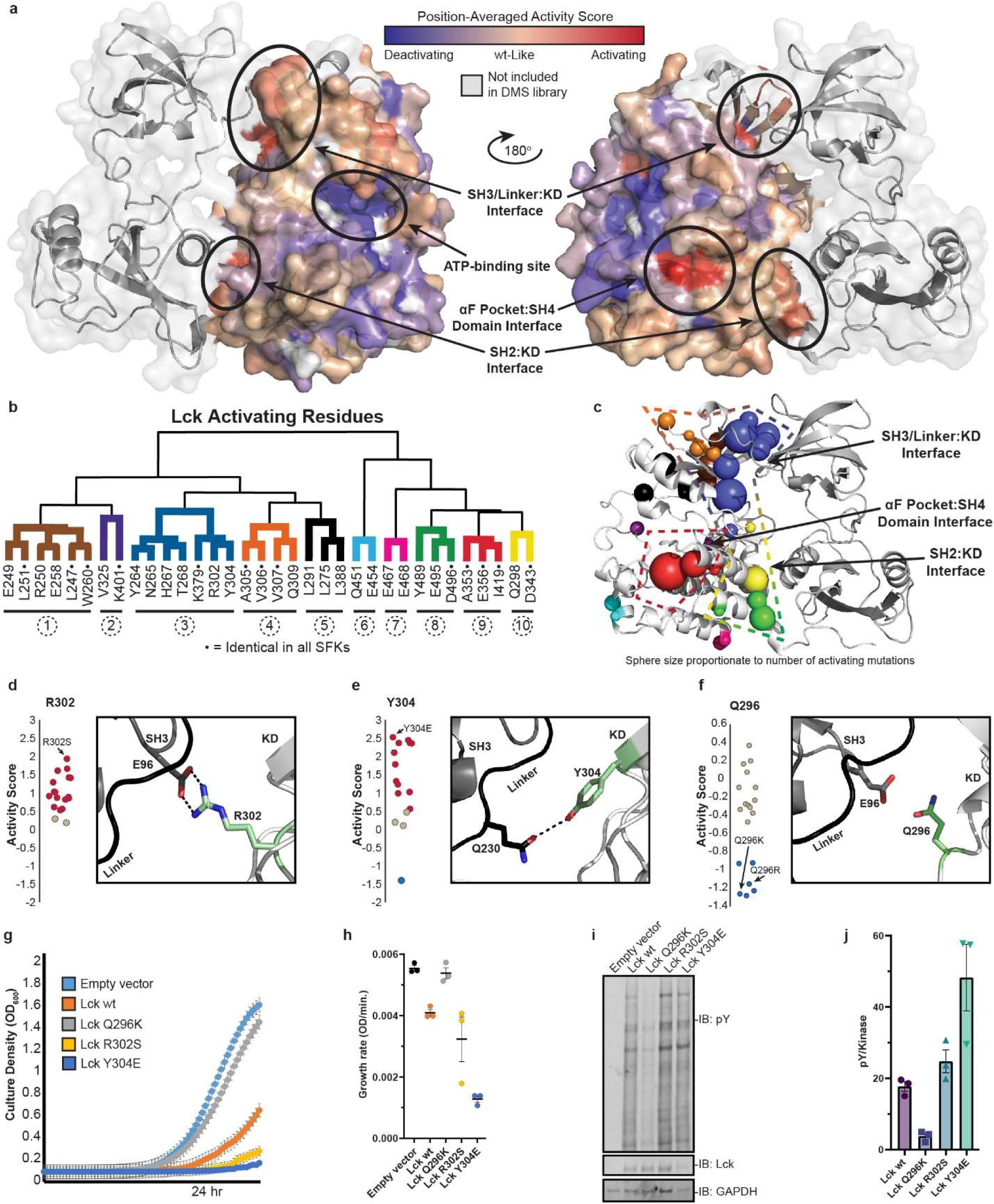
Characterization of Activity-Modulating Variants of Lck. (A) Position-averaged activity scores mapped onto the AlphaFold model of full-length Lck (AlphaFold: AF-P06239-F1; N-terminal SH4 domain, positions 1-72, removed for simplicity). (B) Hierarchical clustering of Lck positions with at least five activating variants based on centroided atomic coordinates for each amino acid side chain extracted from the AlphaFold model of full-length Lck (AlphaFold: AF-P06239-F1). Residues with asterisk as conserved across all Src Family Kinase members. (C) Hierarchical clustering data projected onto the AlphaFold model of Lck (AlphaFold: AF-P06239-F1). Sphere size represents the number of activating mutations per position, colored by clusters from Figure 2B. (D-F) Activity scores for all variants at positions R302, Y304, and Q296 which lay along the SH3/Linker:KD interface. Panels highlight structural detail for corresponding positions and their predicted side chain interactions extracted from the AlphaFold model of full-length Lck (AlphaFold: AF-P06239-F1). (G) Individually assessed growth curves for yeast expressing Lck WT, K273R (kinase dead), R302S, Y304E, Q293K, or a control vector. Datapoints represent average ± s.e.m. of three biological replicates. (H) Growth rates for yeast strains expressing Lck WT, K273R (kinase dead), R302S, Y304E, Q293K, or a control vector. Dots represent individual biological replicates, lines represent averages, error bars represent s.e.m.. (I) Phosphotyrosine immunoblot blot (IB) of lysates from yeast expressing Lck WT, K273R (kinase dead), R302S, Y304E, Q293K, or a control vector. Representative of three replicates. (J) Quantification of phosphotyrosne western blots from Fig. 3I. Dots represent individual biological replicates, bars represent averages, error bars represent s.e.m..

To engineer Lck variants with tuned signaling activity, we focused on three positions at the SH3/linker:KD interface, Q296, R302, and Y304, whose side chains are predicted by AlphaFold to project toward the binding interface formed between the KD, linker, and SH3 domain in the autoinhibited conformation of Lck (**Fig. 3D-3F**). R302 is predicted to form an electrostatic interaction with E96 on the SH3 domain, and most substitutions at this position are activating, consistent with disruption of this contact (**Fig. 3D**). Similarly, the hydroxyl group of Y304 is predicted to be within hydrogen-bonding distance of Q230 on the SH2-KD domain linker, and the majority of substitutions at this position are also activating (**Fig. 3E**). Yeast expressing R302S or Y304E grew more slowly and exhibited elevated phosphotyrosine levels relative to WT, confirming increased phosphotransferase activity (**Fig. 3G-J**). In contrast, Q296, which is predicted to lie in close proximity to E96 in the SH3 domain, is a position where select substitutions appear to enhance autoinhibition. While most substitutions at Q296 demonstrated WT-like Lck activity, Q296K and Q296R were selectively deactivating, consistent with formation of a neo-salt bridge between the introduced basic side chain and E96 that strengthens the autoinhibitory SH3:KD interaction (**Fig. 3F**). Yeast expressing Q296K grew substantially faster than WT and displayed reduced phosphotyrosine levels, with equivalent Lck expression confirmed by immunoblot (**Fig. 3G-J**). Together, these findings establish the SH3/linker:KD interface as a tunable rheostat that permits graded control over Lck phosphotransferase activity through precise manipulation of intramolecular contacts.

### DMS profiling nominates covalent liganding sites that allosterically modulate Lck

We next asked whether deep mutational scanning activity profiles could predict the functional consequences of covalent liganding. We reasoned that the effects of substitutions at solvent-exposed positions occupied by a WT Cys or Lys, and at the positions surrounding them, would report on both the essentiality of the WT position itself and the functional consequences of a covalent ligand occupying an adjacent pocket. We used the PyMOL script FindSurfacePositions to identify all solvent-exposed positions occupied by WT Cys or Lys in a crystal structure of Lck’s KD (PDB 3AD6; Sogabe, et al., 2012). No Cys positions were classified as solvent-exposed, consistent with how poorly these positions tolerate substitution (**Fig. 2C**), whereas 13 Lys positions were (K246, K269, K273, K276, K293, K329, K335, K340, K379, K401, K405, K420, and K478). We projected their position-averaged activity scores onto the AlphaFold model of Lck’s KD (**Fig. 4A**). All substitutions at K273 and K405 were deactivating, demonstrating that the identity of the amino acid at these positions is essential for Lck folding and/or phosphotransferase activity (**Fig. S3A**). The essentiality of K273 is consistent with its established phosphotransferase role in coordinating the α-and β-phosphates of ATP. K405, by contrast, is not known to be involved in catalysis or implicated in the hydrophobic spine model of kinase regulation, though it is located three positions before the APE motif at the C-terminal end of Lck’s activation loop, and is adjacent to positions that form the hydrophobic spine. Therefore, we speculate that perturbations to K405, via amino acid substitution or chemical modification, inhibit Lck activity by acting on one or more of these regulatory elements through an allosteric mechanism.

**Figure 4.**
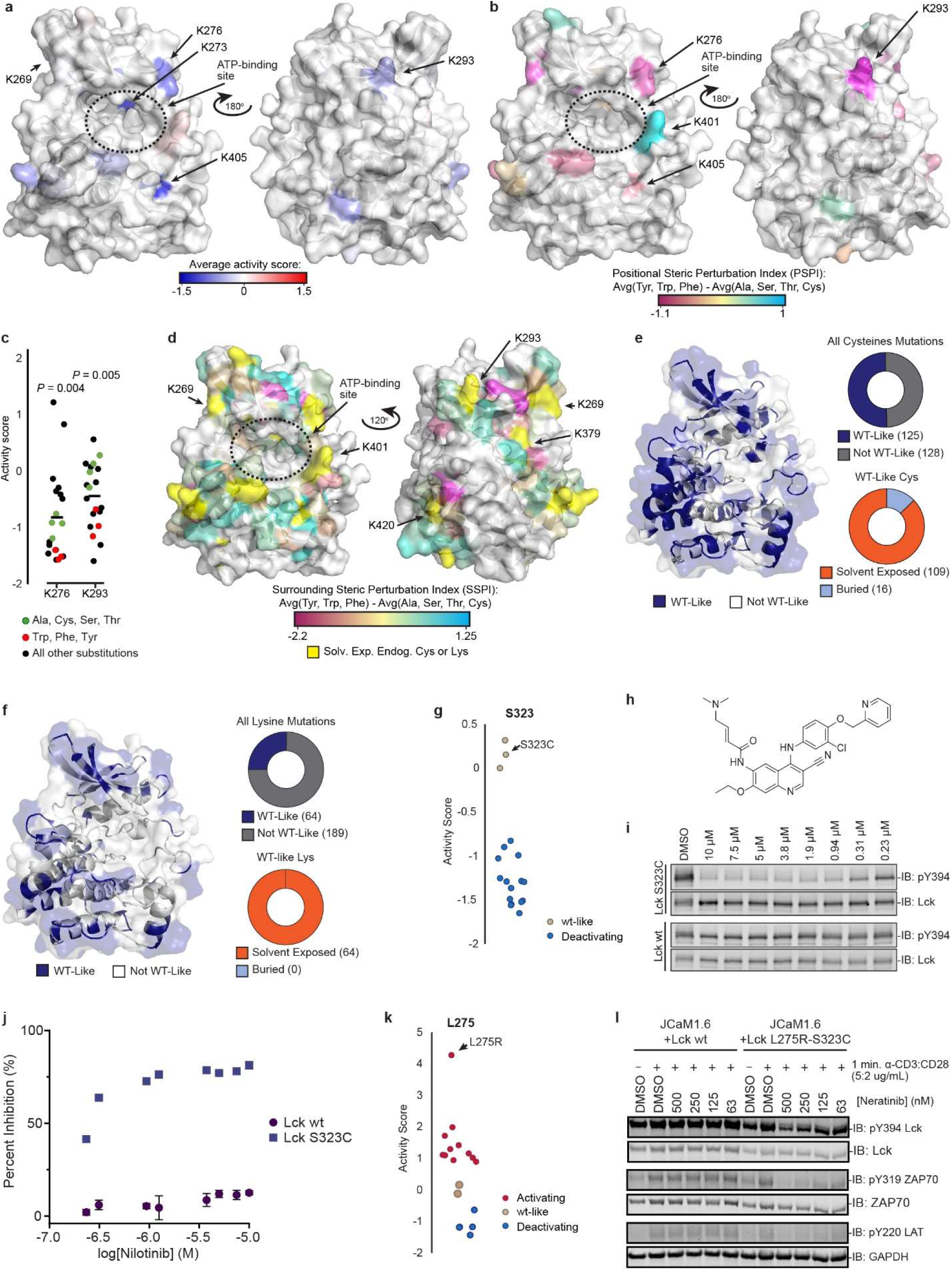
Analysis of mutational sensitivity near endogenous Lys amino acids and WT-like Cys and Lys amino acid substitutions. (A) Position-averaged activity scores for solvent-accessible endogenous lysine positions mapped onto AlphaFold model of Lck’s KD (AlphaFold: AF-P06239-F1). (B) Positional Steric Perturbation Index (PSPI) scores for solvent exposed endogenous Lys positions mapped onto the AlphaFold model of Lck’s KD (AlphaFold: AF-P06239-F1). Scores are derived for each neighboring position by averaging the activity score of Tyr, Phe, and Trp, and subtracting the average activity score of Ala, Ser, Thr, and Cys. (C) Activity scores for every variant for positions with statistically significant differences between average of Phe, Tyr, and Trp activity scores and average of Ala, Gly, Ser, Thr, and Cys activity scores. p-values were calculated using a two-tailed Student’s t test. (D) Surrounding Steric Perturbation Index (SSPI) scores for amino acids within 5 Å of endogenous Lys positions mapped onto the AlphaFold model of Lck’s KD (AlphaFold: AF-P06239-F1). Scores are derived for each neighboring position by averaging the activity score of Tyr, Phe, and Trp, and subtracting the average activity score of Ala, Gly, Ser, Thr, and Cys, and then averaged to yield a single SSPI score for each endogenous Lys position. (E) (left) WT-like cysteine positions (blue) mapped to the AlphaFold AlphaFold model of Lck’s KD (AlphaFold: AF-P06239-F1). (top right) Pie chart represents all WT-like cysteine mutations (blue) on Lck’s KD. (bottom right) Pie chart represents all solvent exposed WT-like cysteine mutations (orange). (F) (left) WT-like lysine positions (blue) mapped to the AlphaFold AlphaFold model of Lck’s KD (AlphaFold: AF-P06239-F1). (top right) Pie chart represents all WT-like lysine mutations (blue) on Lck’s KD. (bottom right) Pie chart represents all solvent exposed WT-like lysine mutations (orange). (G) Activity scores for all S323 variants; S323C highlighted with an arrow. (H) Chemical structure of neratinib. (I) Immunoblot analysis (IB) of HEK293T cells transiently expressing WT or S323C Lck treated with neratinib. Cells expressing WT or S323C Lck were treated with various concentrations of neratinib for 4 h, lysed and subjected to SDS-PAGE and immunoblot analysis. Representative of two independent experiments. (J) Quantification of pY394 inhibition from experiments shown in 4I. (K) Activity scores for all L275 variants; L275R highlighted with an arrow. (L) Immunoblot analysis (IB) of JCaM1.6 cells stably expressing WT or S323C-L275R Lck pre-treated with indicated concentration of neratinib (1 h) prior to TCR activation with anti-CD3:CD28 antibody cocktail (5:2 µg/mL; 1 min). Cells were then quenched on ice, lysed and subjected to SDS-PAGE and immunoblot analysis.

Identifying positions intolerant of substitution reveals amino acids important for catalysis and/or folding, where covalent modification would likely inactivate Lck, but it does not directly predict the functional consequence of a ligand occupying an adjacent binding pocket. We therefore developed a Steric Perturbation Index (SPI) to model this scenario, reasoning that bulky aromatic substitutions (Phe, Trp, and Tyr) best approximate the steric footprint of a covalent ligand engaging a pocket adjacent to a nucleophilic amino acid, while more modest substitutions (Ala, Thr, Ser, and Cys) report on the functional consequences of losing the WT amino acid side chain itself. Positional SPI values (PSPI) were calculated as the difference between the average activity scores of these two substitution classes at each solvent-exposed Lys position (**Fig. S3A** and **S3B**, middle column) and projected onto the AlphaFold model of Lck’s KD (**Fig. 4B**), with a negative or positive PSPI predicting that covalent engagement at that site would inhibit or activate kinase activity, respectively. We first applied this reasoning to the solvent-exposed Lys positions themselves. Two positions, K276 and K293, had statistically significant negative PSPI scores, nominating them as candidate sites where chemoselective covalent liganding is predicted to allosterically inhibit Lck phosphotransferase activity (**Fig. 4B, 4C,** and **S3A**).

We next analyzed the substitutions at positions surrounding each solvent-exposed endogenous Lys amino acid to identify putative binding sites that could potentially alter Lck’s activity upon successful liganding. We calculated SPI scores for every position within 5 Å of each solvent-exposed endogenous Lys, averaged these scores to give surrounding SPI (SSPI) scores (**Fig. S3B**, right) and projected them onto the AlphaFold model of Lck’s KD (**Fig. 4D**). K379, K401, and K420 had the most negative SSPI scores, suggesting that liganding at these sites could result in inhibition (**Fig. S3C**). Notably, none of these three Lys positions are themselves implicated in the hydrophobic spine, involved in catalysis, or located within 5 Å of the ATP-binding site, so their strongly negative neighborhoods cannot be explained by direct interference with the catalytic machinery. K420 is located in the C-lobe, and its side chain makes hydrogen bonds with the backbone amides of a neighboring loop, likely providing structural stability. Two of these positions, K379 and K401, each harbor 7 activating substitutions and were therefore identified in our hierarchical clustering analysis (**Fig. 3B**, clusters 3 and 2, respectively), and both are conserved across all SFKs. These data echo the pattern we noted above for K405, where perturbation of a site outside the catalytic core nonetheless carries functional consequences on phosphotransferase activity. More generally, we speculate that the SSPI scores at these positions reflect allosteric coupling between these regions and the ATP-binding site, a bidirectional relationship that can activate or deactivate the kinase depending on the nature of the perturbation. For example, large aromatic substitutions at K401, which resides on one face of a groove on the C-lobe, were strongly activating (**Fig. 4B** and **S3A**), whereas substitutions at the neighboring K405 on the opposing face were uniformly deactivating (**Fig. 4A, 4B**, and **S3A)**, suggesting that covalent engagement of adjacent sites within the same surface pocket could produce opposing allosteric outcomes. Together, these analyses nominate specific endogenous Lys positions and their adjacent pockets as candidate sites for covalent small-molecule targeting and highlight how deep mutational scanning can reveal cryptic opportunities for chemical control over protein function.

### WT-like cysteine and lysine substitutions provide handles for ligand sensitization

We reasoned that positions where Cys or Lys substitutions preserve WT-like activity, and that lie near canonical or cryptic small-molecule binding pockets, could be leveraged to sensitize Lck to electrophile-containing small molecules. To identify such sites, we projected all KD positions that maintained WT-like activity when substituted with Cys or Lys onto the structure of Lck’s KD (**Fig. 4E** and **4F**). These positions are widely distributed across Lck’s KD, with WT-like Cys substitutions covering 50.5% of Lck’s solvent-exposed surface area and WT-like Lys substitutions covering 37.1%. Almost all of these positions are solvent-exposed, suggesting many opportunities for covalent targeting.

To demonstrate how introduced Cys or Lys substitutions can be leveraged to confer ligand sensitivity, we evaluated WT-like Cys substitutions near the ATP-binding site for their potential to sensitize Lck to FDA-approved electrophile-containing kinase inhibitors. Structural alignment of the Lck and EGFR KDs revealed that EGFR C797, which forms a covalent adduct with multiple FDA-approved EGFR inhibitors, aligns with S323 in Lck (**Fig. S3D**). S323C was classified as having WT-like activity (**Fig. 4G** and **1B**), suggesting that a Cys substitution at this position could sensitize Lck to covalent EGFR inhibitors, such as neratinib (**Fig. 4H**), while preserving phosphotransferase function. S323C Lck also underwent levels of activation loop autophosphorylation comparable to WT Lck when transiently expressed in HEK293T cells (**Fig. 4I**), confirming that its phosphotransferase activity is conserved in a human cell environment.

Although most clinically approved covalent EGFR inhibitors are highly selective and minimally target Lck, a comprehensive chemoproteomic profiling study (Klaeger, et al., 2017) identified Lck as a weak off-target of neratinib. We therefore tested whether introducing Cys at position 323 could convert the weak non-covalent targeting of WT Lck into potent inhibition of S323C Lck in human cells. As expected, neratinib minimally inhibited WT Lck activation loop autophosphorylation at all concentrations tested, whereas S323C Lck activation loop autophosphorylation was potently inhibited in a dose-dependent manner (IC_50_ < 300 nM; **Fig. 4I** and **4J**). We also combined the S323C substitution with the activating L275R substitution (L275R-S323C; **Fig. 4K**), and measured blockade of TCR activation-dependent phosphorylation events in Lck-null JCaM1.6 cells recombinantly expressing WT or L275R-S323C Lck. Although neratinib pre-treatment did not prevent the phosphorylation of ZAP70 (pY319) and LAT (pY220) in WT Lck-expressing JCaM1.6 cells, both events were potently blocked in cells expressing L275R-S323C Lck (**Fig. 4L**). Together, these results demonstrate the ability to rationally select covalent kinase inhibitor/Lck variant pairs, providing a foundation for engineering drug-sensitive Lck variants with potential applications in T cell-derived immunotherapeutics.

### Identification of Lck variants sensitized to reversible ATP-competitive inhibitors

Having identified substitutions that confer sensitivity to covalent inhibition, we next sought to determine whether Lck could be sensitized to reversible ATP-competitive inhibitors. To maximize translational relevance and potential for *in vivo* use, we restricted our search to clinically approved drugs that have favorable long-term safety profiles and otherwise limited off-target kinase activity, but that retain weak affinity for Lck. This nominated the EGFR inhibitors erlotinib and gefitinib, the BRAF inhibitor dabrafenib, and the BCR-ABL inhibitors nilotinib and imatinib, all of which have weak off-target activity against Lck (Klaeger, et al., 2017; Médard, et al., 2015; Davis, et al., 2011; Karaman, et al., 2008), and have been chronically administered to patients with manageable toxicity profiles (Shepherd, et al., 2005; Hauschild, et al., 2012; Kris, et al., 2003; Kantarjian, et al., 2011). Notably, these inhibitors stabilize distinct ATP-binding site conformations, raising the possibility that they could differentially modulate regulatory domain accessibility, and thus scaffolding function, of a sensitized Lck variant (Fang, et al., 2020). Imatinib and nilotinib stabilize their target Abl in a DFG-out, αC-helix-in conformation, which in SFKs promotes disengagement of the regulatory domains and engagement of signaling partners. Dabrafenib stabilizes its target BRAF in a DFG-in, αC-helix-out conformation, which in SFKs promotes further intramolecular engagement of the regulatory domains. Erlotinib and gefitinib minimally alter the conformation of EGFR’s ATP-binding site. Together, these drugs span a range of ATP-site conformations, providing a clinically approved toolkit for targeting sensitized Lck variants.

To systematically identify Lck variants with enhanced sensitivity to small-molecule inhibition, we used the yeast-based growth assay to measure how well each drug rescued the growth inhibition caused by Lck phosphotransferase activity. Because small-molecule screening in yeast is often limited by rapid efflux, we performed all Lck variant profiling in a genetically engineered ABC transporter deletion strain (ABC16; Suzuki, et al., 2011). To confirm that this strain was appropriate for our drug sensitization efforts, we first verified that nilotinib, one of the drugs selected for sensitization screening, increased the growth of this strain when it expressed WT Lck, but not when it expressed a catalytically inactive (K273R) variant **(Fig. S4A**). We then identified the lowest concentration of each drug that produced detectable inhibition of WT Lck, defined as a ∼10% reduction in bulk phosphotyrosine, so that sensitized variants could be identified under conditions where WT Lck remained largely uninhibited (**Fig. S4B-S4D**).

The Lck variant library was transformed into yeast, and a sample was collected prior to Lck induction. Lck expression was then induced in the presence of either DMSO or each drug at the concentration that gave minimal but detectable inhibition of WT Lck (**Fig. S4C** and **S4D**), and samples were collected throughout outgrowth. Plasmids were extracted, and activity scores for each inhibitor were calculated from two independent biological replicates (r between 0.82 and 0.88; **Fig. S4E**) as described above for the activity profiling experiments. For each variant, we then calculated a “sensitization score” as its drug-treated activity score minus its DMSO activity score, such that more negative scores indicate greater drug sensitization (**Fig. 5A**). Sensitization scores were obtained for an average of 4,871 variants per drug. As expected, most variants, including synonymous WT variants, had sensitization scores near zero, indicating either that they were not sensitized to the drug tested or that they lacked sufficient activity for a difference in inhibition to be measured (**Fig. 5A** and **S4F**). To validate these measured sensitization scores, we expressed four variants spanning a range of imatinib sensitization scores, alongside WT and catalytically inactive (K273R) Lck, in yeast and measured growth in the presence of DMSO or imatinib (12.5 µM). Growth correlated well with the measured imatinib sensitization scores (r² = 0.9; **Fig. 5B** and **5C**).

**Figure 5.**
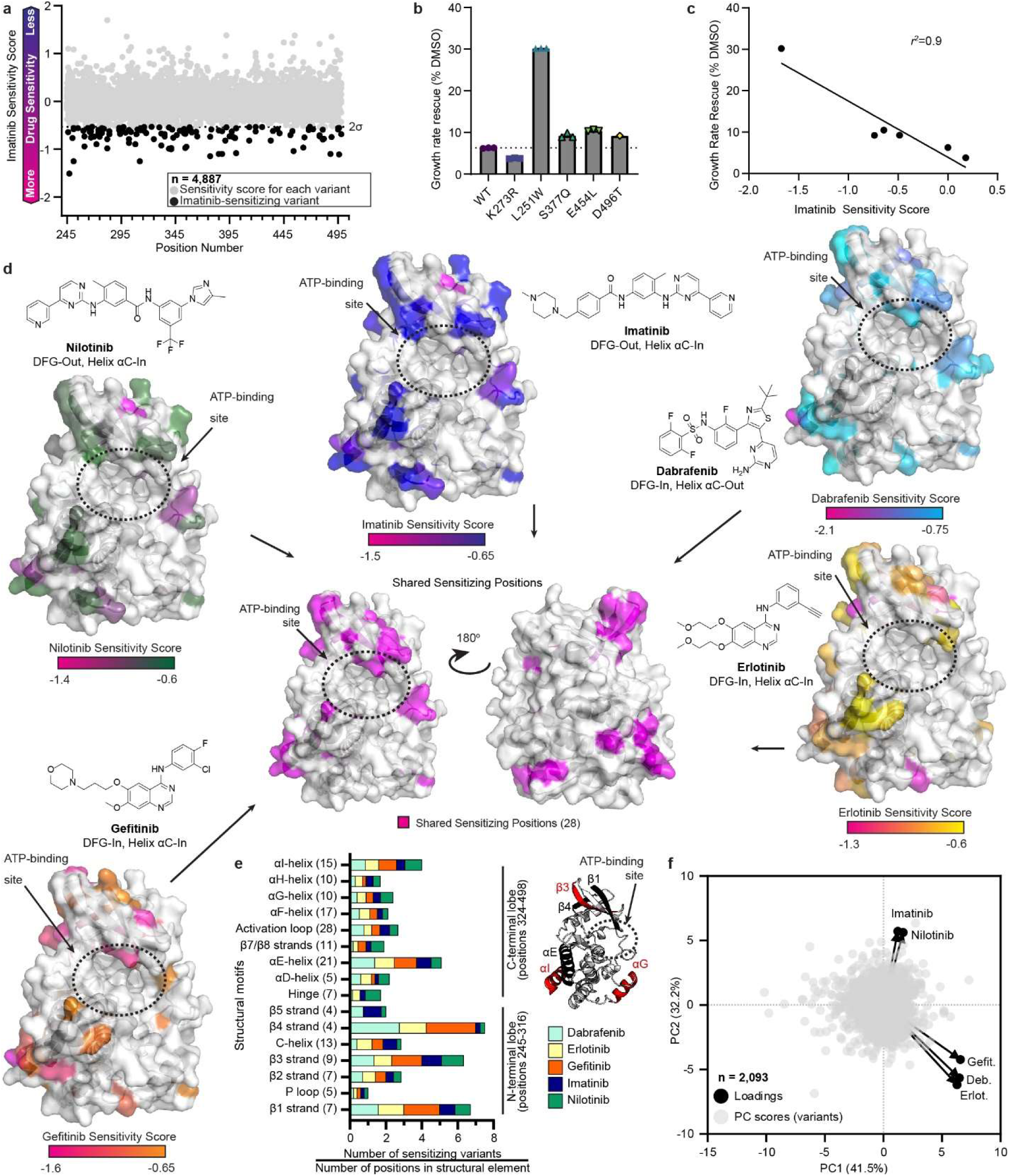
Screen of 4,975 Single Amino Acid Lck Variants for Drug Sensitization. (A) Sensitivity scores for all variants quantified in the imatinib sensitization screen. Dotted line represents two standard deviations from the mean sensitization score. Sensitization events are those variants with scores below this line. (B) Growth rates for yeast strains expressing Lck WT, K273R (kinase dead), L251W, S377Q, E454L, or D496T grown in the presence of 12.5 µM imatinib relative to those grown in DMSO. Growth rate rescue calculated as a percentage of growth rate in the presence of DMSO compared to inhibitor treatment. Dots represent individual biological replicates, bars represent averages, error bars represent s.e.m.. (C) Correlation between imatinib sensitivity scores and growth rate rescue of individual variants grown in the presence of 12.5 µM imatinib measured in 5B. (D) Position-averaged sensitization score for each inhibitor mapped onto the AlphaFold model of Lck’s KD (AlphaFold: AF-P06239-F1). (center) Shared sensitizing positions (28), defined as amino acid positions with at least one sensitization event per drug mapped onto the AlphaFold model of Lck’s KD. Number of sensitizing variants per inhibitor: Dabrafenib, 189; Erlotinib, 164; Gefitinib, 176; Imatinib, 130; Nilotinib, 138. Number of sensitizing positions per inhibitor: Dabrafenib, 89; Erlotinib, 91; Gefitinib, 87; Imatinib, 92; Nilotinib, 94. (E) left, Sensitizing events stratified by Lck’s structural motifs. Bar represents all sensitization events for a given structural motif divided by the number of positions in that motif. The shaded region of each bar represents the proportion of the total number of variants on that motif belonging to each inhibitor. right, AlphaFold model of Lck’s KD (AlphaFold: AF-P06239-F1) with prominent structural motifs shaded in red or black. (F) Principal component analysis of inhibitor sensitivity scores.

We defined drug-sensitized variants, or “sensitization events,” as those with sensitization scores more than two standard deviations (2σ) below the mean for each drug, and we mapped their sensitization scores onto the structure of the Lck KD (**Fig. 5D**). This classification yielded 130 to 189 sensitization events per drug, an average of 63 of which were unique to a single drug. These sensitization events occurred at an average of 91 positions per drug, 5 to 15 of which were unique to a single drug, corresponding to an average of 1.4 (imatinib) to 2.1 (dabrafenib) sensitization events per position (**Table 1**).

**Table 1.**
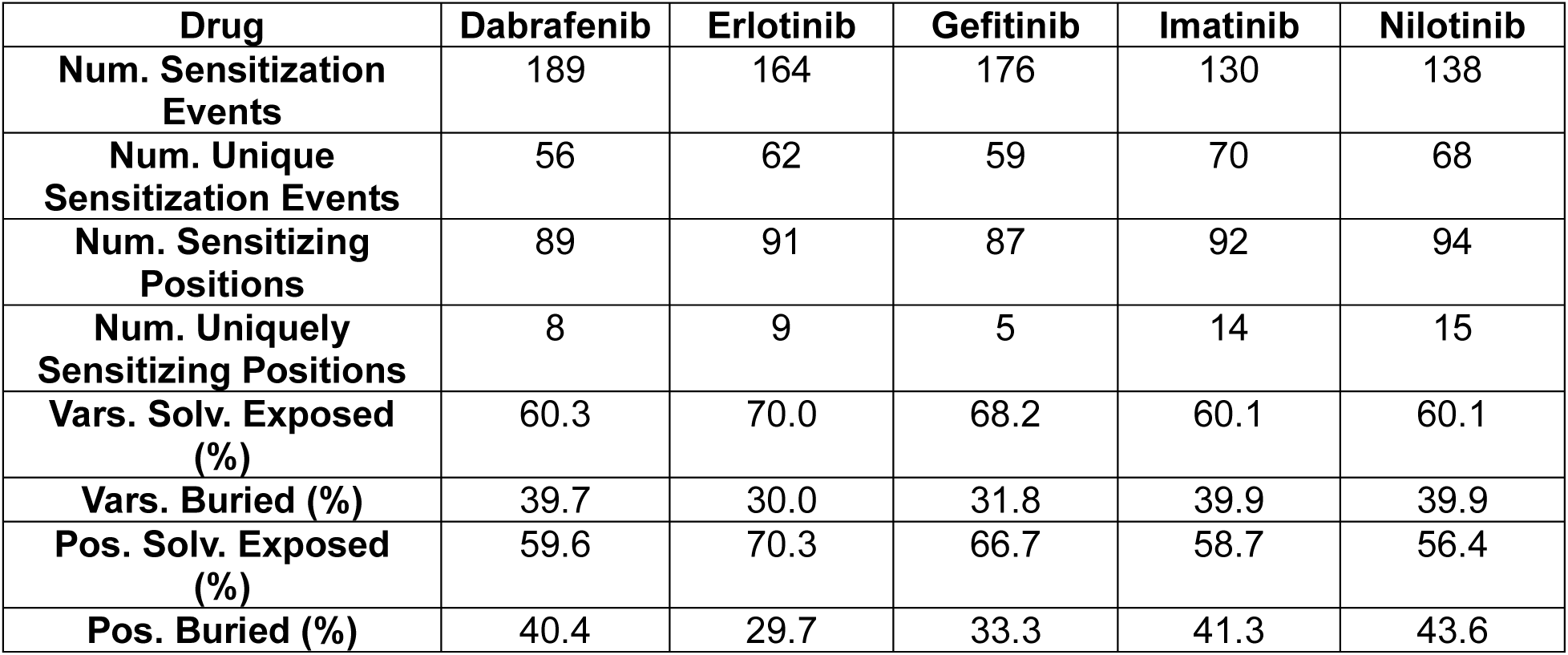
Number of drug-sensitized variants and positions for each inhibitor dataset.

### Sensitization arises through long-range allosteric mechanisms

Having identified 797 sensitizing events (**Table 1**, top row), we next asked which features distinguish sensitization events and where in Lck they occur. After normalizing for the frequency of each amino acid in Lck, substitutions at positions occupied by Gln and Glu were the most likely to yield a sensitization event, while those at positions occupied by Cys and Gly were the least likely (**Fig. S4G**). Substitution to Gln and Ser, followed by Ile and Thr, contributed the largest percentage of sensitization events, while substitution to Pro and Arg contributed the smallest (**Fig. S4G**). Sensitization events were also enriched at positions with a relative solvent-exposed surface area (rSASA) > 0.2 Å^2^ (64% versus 51% of all positions, 0.34 rSASA versus 0.29; p < 0.01; **Fig. S4H**), suggesting that solvent-exposed positions more readily support drug sensitization. To determine whether sensitization arises mainly through alteration of local drug interactions in the ATP-binding site or through more indirect allosteric mechanisms, we calculated the distance from the center of mass of imatinib bound to Lck (PDB 2PL0; Jacobs, et al., 2008) to the α-carbon of each position on Lck’s KD. Sensitization events were located a median distance of 17.7 Å from imatinib’s center of mass (with 25% of variants >24.2 Å), suggesting that many variants confer drug sensitivity through indirect allosteric mechanisms (**Fig. S4I**). Of all sensitizing events, 71.1% resided within well-defined kinase structural motifs, with the highest density within the N-lobe (**Fig. 5E**). Notably, the β1 (positions 245-251), β3 (positions 267-275), and β4 (positions 304-307) strands contributed the most sensitizing events per position, and a total of seven positions on these three strands harbored sensitization events for all five drugs, though not necessarily the same variant in each case. In the C-lobe, the αE-and αI-helices harbored the largest number of sensitizing events on a per-position basis. The remaining 28.9% were located on loops connecting structural motifs (**Fig. S4J**). Among these, the β2/β3-strand loop (positions 264-266) was the most frequently sensitized loop, harboring an average of 11.7 sensitizing events per position. Together, these analyses indicate that sensitization events are broadly distributed across the KD, favoring solvent-exposed positions remote from the ATP-binding site, and that they are enriched on several β strands on the N-lobe.

We next asked whether any positions in Lck conferred sensitization to all five drugs when varied. We found 28 positions that had at least one sensitization event for every drug (315 variants in total; **Fig. 5D**, center; **Fig. S4K**). Similar to our analysis of all sensitization events, we found that these 28 positions were located an average of 19.7 Å from imatinib’s center of mass (PDB 2PL0; Jacobs, et al., 2008), indicating that they too mostly function by communicating allosterically with the ATP-binding site. Although 28 positions had sensitization events for every drug, the identity of the sensitization event differed between drugs; only two individual variants, L388S and Q451L, were sensitized to all five drugs. Position L388 lies on the activation loop, four positions C-terminal to the DFG-motif, and introducing a Ser into this conformationally flexible region appears to allow all five inhibitors to be better accommodated. Q451 lies on the solvent-exposed face of the αG-helix, 32 Å from the ATP-binding site. We have previously shown that this region is allosterically coupled to the ATP-binding site in Src (Potter, et al., 2020), and replacing Gln with a hydrophobic Leu at this position appears to transmit a conformational change that sensitizes the ATP-binding site to all five drugs. Of the 28 shared sensitizing positions, 23 were located on kinase structural motifs, with the β3 strand having the highest percentage (4 of 9) of positions capable of providing sensitization to all five drugs when varied. Of the remaining five positions not on kinase structural motifs, two were located on the β2/β3 loop, indicating that the β3 strand region provides the most general route to drug sensitization when varied. Together, these analyses suggest that inhibitor sensitization arises through long-range allosteric mechanisms rather than the direct modulation of inhibitor contacts.

### Substitution identity and inhibitor class encode distinct conformational signatures

Specific substitutions at individual positions might facilitate the conformational preferences of the three drug classes we tested. To explore this possibility, we focused first on R302, which harbored the most sensitization events (26) among the 28 positions sensitized to all five drugs (**Fig. S4L**). Substitutions at this position partitioned by inhibitor class: nine variants involved dabrafenib, four of which (R302V, R302L, R302Y, and R302C) were also sensitized to erlotinib and gefitinib, whereas imatinib and nilotinib shared two sensitization events with each other (R302G and R302D) that did not overlap with those of dabrafenib, erlotinib, or gefitinib. R302 sits within the SH3/linker:KD interface (**Fig. 3B**, **3C**), and many substitutions at this position increase Lck phosphotransferase activity (**Fig. 3D**), so sensitization here is likely to arise both from direct allosteric modulation of the ATP-binding site conformation and from altered regulatory domain interactions. The positions with the next largest numbers of sensitization events each harbored 19 and displayed similarly class-dependent patterns of sensitization: T268, on the β3-strand is also part of the SH3/linker:KD interface, and D343, on the αE-helix within the SH2:KD interface (**Fig. S4L**). Together, these results suggest that although DFG-out and DFG-in inhibitors converge on many of the same 28 positions, they achieve sensitization through at least partially distinct structural mechanisms, especially at positions forming regulatory domain interfaces.

Aggregating sensitizing positions by inhibitor class offered a second route to unveiling class-specific conformational preferences. We identified every position harboring at least one sensitization event for a given inhibitor, which allowed us to separate positions uniquely sensitizing to a single inhibitor from those shared across inhibitors (**Fig. S4K**). Each inhibitor had an average of 10 uniquely sensitizing positions, the two DFG-out inhibitors (imatinib and nilotinib) shared eight positions, and the three DFG-in inhibitors (dabrafenib, erlotinib, and gefitinib) shared 10. Measuring the average distance of these class-specific positions from imatinib’s center of mass (PDB 2PL0; Jacobs, et al., 2008) revealed that DFG-out-sensitizing positions lie closer to the ATP-binding site and the DFG-motif than DFG-in-sensitizing positions (14.0 Å versus 20.3 Å; p = 0.02; **Fig. S4M**). Sensitization to the two inhibitor classes is therefore encoded at different distances from the site of drug binding.

### Principal component analysis of sensitization scores segregates inhibitors by ATP-binding site conformation

To capture the conformational signatures encoded across all five sensitization datasets, we applied principal component analysis (PCA) to the sensitization score matrix. The first two principal components captured 73.7% of the total variance (PC1, 41.5%; PC2, 32.2%), and projection of the inhibitor loadings grouped the compounds according to their known conformational preferences (**Fig. 5F**). The two DFG-out-stabilizing inhibitors, imatinib and nilotinib, co-segregated along PC2, while erlotinib and gefitinib, which stabilize the opposing DFG-in conformation, clustered in the opposite direction, as predicted from a DFG-motif-based stratification. Dabrafenib provided a more informative test of this model. Although it stabilizes a DFG-in/αC-out conformation distinct from the DFG-in/αC-in state stabilized by erlotinib and gefitinib, dabrafenib co-clustered with those two inhibitors along PC2, resolved from imatinib and nilotinib, indicating that, at least within this inhibitor set, DFG-motif orientation is the dominant conformational signature captured by sensitization scores, outweighing any contribution from αC-helix state. Together with the lobe-and secondary-structure-level distributions described above, these results support a model in which inhibitor sensitization is governed by a distributed allosteric network that principally senses and communicates with the conformational state of the DFG-motif.

### Biochemical validation of nilotinib-sensitized Lck variants

We next asked whether the increased inhibitor sensitivity observed in yeast would translate to human cells. To test this, we focused on 26 of the most nilotinib-sensitizing variants (**Fig. 6A**, black dots). We ectopically expressed WT Lck and the 26 candidate drug-sensitized variants in HEK293T cells, treated them with 5 µM nilotinib for 4 hours, and measured activation loop phosphorylation levels. As expected, WT Lck showed a minimal reduction in activation loop phosphorylation; however, more than half of the candidate drug-sensitized variants exhibited enhanced inhibition of activation loop phosphorylation (performers; **Fig. 6B** and **S5A**). Dose-response analysis revealed that single substitutions (e.g., L251W, I312S) can increase nilotinib sensitivity by approximately 10-30-fold (**Fig. 6C** and **S5B**). Together, these results confirm that yeast-based sensitization scores predict nilotinib sensitivity in human cells, nominating a set of candidate variants for functional validation in T cells.

**Figure 6.**
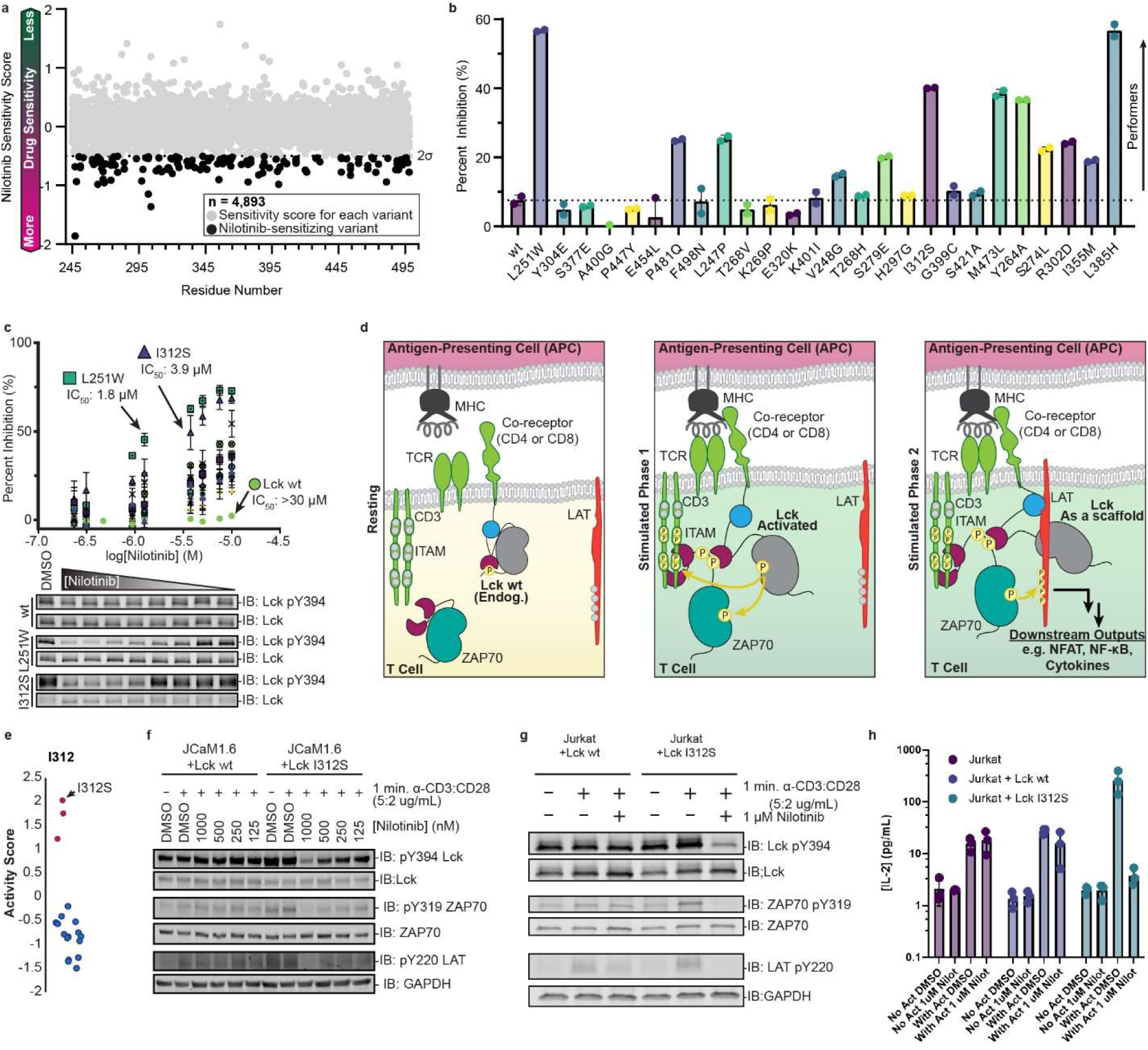
Lck I312S demonstrates sensitivity to nilotinib, increases antigen sensitivity in JCaM1.6 cells, acts as an off switch in T Cell activation assays. (A) Sensitivity scores for all variants quantified in the nilotinib sensitization screen. Dotted line represents two standard deviations from the mean sensitization score. Sensitization events are those variants with scores below this line. (B) Quantification of immunoblots (IB) measuring Lck activation loop phosphorylation levels (pY394) for nilotinib-sensitive variants identified in yeast, ectopically expressed in HEK293T cells and treated with DMSO or 5 µM nilotinib for 4 h. Dots represent individual biological replicates, bars represent averages (n=2). (C) top. Quantification of immunoblots (IB) measuring activation loop phosphorylation levels of nilotinib-sensitive variants transiently expressed in HEK293T cells and treated with various concentrations of nilotinib for 4 h (n=2). bottom. Representative immunoblots (IB) for WT, L251W, or I312S Lck ectopically expressed in HEK293T cells treated with various concentrations of nilotinib for 4 h. Representative to two independent biological replicates. (D) Schematic of T cell activation highlighting Lck’s role as an initiator and scaffold. Adapted from Lo, et al., 2018. (E) Activity scores for all I312 variants; I312S highlighted with an arrow. (F) Immunoblot analysis (IB) of JCaM1.6 cells recombinantly expressing WT or I312SLck pre-treated with the indicated concentration of nilotinib (1 h). Cells were then stimulated with anti-CD3:CD28 antibody cocktail (5:2 µg/mL, 1 min), quenched on ice, lysed and subjected to SDS-PAGE and immunoblot analysis. Representative of two independent experiments. (G) Immunoblot analysis of parental Jurkat cells or those recombinantly expressing WT or I312S Lck pre-treated with or without nilotinib (1 µM, 1 h). Cells were then stimulated with anti-CD3:CD28 antibodies (5:2 µg/mL, 1 min), quenched on ice, lysed and subjected to SDS-PAGE and immunoblot analysis. Representative of two independent experiments. (H) IL-2 levels from parental Jurkat cells, or Jurkat cells recombinantly expressing WT or I312S pre-treated with DMSO or nilotinib (1 µM, 1 h), followed by stimulation with anti-CD3:CD28 antibody cocktail (5:2 µg/mL, 6 hr). After quenching, IL-2 levels were measured using an ELISA. Dots represent individual biological replicates. Bars represent an average of three replicates ± s.e.m.

### Drug-sensitized Lck variants enable drug-gated control over T cell signaling outputs

We next sought to validate inhibitor-sensitized Lck variants in a more physiologically relevant setting. Upon TCR engagement, Lck initiates signaling by phosphorylating immunoreceptor tyrosine-based activation motifs (ITAMs) within the CD3-ζ chains, creating docking sites for ZAP70 (**Fig. 6D**). Lck subsequently phosphorylates and activates ZAP70 while also functioning as a scaffold that promotes productive interactions between ZAP70 and its substrate, LAT, at the plasma membrane (**Fig. 6D**). This coordinated phosphotransferase and scaffolding activity enables efficient LAT phosphorylation, assembly of the LAT signalosome, and propagation of the downstream signaling required for T cell activation (Lo, et al., 2018). We reasoned that an inhibitor-sensitized Lck variant stabilized in the open global conformation by a DFG-out, αC-helix-in inhibitor (e.g., nilotinib or imatinib) could allow an sensitized variant engaged by an inhibitor to outcompete endogenous WT Lck for signaling partners. The I312S variant emerged as a compelling candidate, being the most nilotinib-sensitive variant tested (IC_50_ of 3.9 µM in HEK293T cells; **Fig. 6B, S5A**, and **S5B**) that was not hypomorphic (compare L251W activation loop phosphorylation levels to WT in **S5A**). In fact, I312S was classified as activating in our activity profiling experiment, a result later confirmed in HEK293T cells (**Fig. 6E** and **S5A**). Its combination of high nilotinib sensitivity and preserved phosphotransferase activity made I312S an excellent candidate for testing drug-gated control of Lck-dependent signaling in T cells.

We investigated the signaling properties and tunability of the I312S variant first in the Lck deficient JCaM1.6 cell line. Consistent with its greater phosphotransferase activity in yeast and HEK293T cells, I312S Lck promoted greater basal activation loop phosphorylation (pY394) and downstream signaling through ZAP70 (pY319) and LAT (pY220) than WT Lck (**Fig. 6F** and **S5C**). Pre-treatment with 1 µM nilotinib nearly completely suppressed Lck activation loop phosphorylation as well as pY319 ZAP70 and pY220 LAT in I312S-expressing cells, with no observable effect in cells expressing WT Lck, confirming that the inhibitory effects of nilotinib act specifically through the sensitized I312S variant (**Fig. 6F** and **S5C**). A clinically relevant implementation of an engineered Lck variant would likely involve viral integration in autologous or allogeneic T cells, where endogenous Lck, left unsensitized to nilotinib, could permit residual signaling flux even in the presence of inhibitor. We therefore asked how I312S Lck would behave in Jurkat cells, which express endogenous Lck, alongside the engineered I312S variant. Consistent with our observations in JCaM1.6 cells, I312S Lck increased activation loop phosphorylation (pY394) and downstream signaling markers pY319 ZAP70 and pY220 LAT (**Fig. 6G** and **S5D**), and 1 µM nilotinib pre-treatment selectively inhibited TCR activation-dependent signaling in I312S Lck-expressing Jurkat cells relative to those expressing WT Lck (**Fig. 6G** and **S5D**). These results demonstrate that I312S confers nilotinib-gated control over proximal TCR signaling even in the presence of endogenous, drug-insensitive WT Lck.

Encouraged by these results, we compared IL-2 cytokine production from parental Jurkat cells and those expressing either WT or I312S Lck following nilotinib pre-treatment (**Fig. 6H**). Nilotinib had no effect on IL-2 production in stimulated parental Jurkat cells or those expressing WT Lck; however, nilotinib treatment prevented IL-2 production in I312S Lck-expressing cells (**Fig. 6H**). This result extends drug-gated control from proximal signaling events (Lck activation loop, ZAP70, and LAT phosphorylation) to a downstream, functionally meaningful T cell output, demonstrating that the I312S Lck/nilotinib pair can govern a physiologically relevant endpoint of T cell activation.

## Discussion

Precise chemical control over cell signaling is a central goal in both basic biology and in the engineering of cell-derived therapies. Here, we establish a framework for engineering small molecule-controlled signaling enzymes by systematically mapping the effects of single amino acid substitutions on enzymatic activity and small molecule inhibition. Through deep mutational scanning of the multidomain tyrosine kinase Lck in the presence and absence of small molecule inhibitors, we demonstrate that such enzymes can be sensitized to FDA-approved covalent and reversible inhibitors while preserving or even enhancing their signaling function. These findings extend chemical-genetic strategies beyond classical active site engineering efforts and provide a generalizable route to programmable control of cellular signaling outputs.

An important insight from this work is that small molecule sensitivity appears to be a property that is distributed throughout a protein, rather than being confined to positions that are known to directly contact small molecules. Specifically shown here, inhibitor-sensitizing substitutions were identified throughout Lck’s KD, including at positions more than 30 Å from the ATP-binding site, implicating long-range allosteric networks in governing ATP-competitive inhibitor binding. Our data further suggest that even single amino acid substitutions can lower the energetic barrier to stabilizing inhibitor-compatible ATP-binding site conformations, such as the “DFG-out, αC-in” state favored by nilotinib and imatinib, thereby increasing binding affinity without introducing new direct contacts with the inhibitor itself. These results underscore the power of empirical mutational mapping in revealing allosteric networks that are difficult to predict from sequence or structure alone.

Our analysis of deactivating substitutions proximal to endogenous lysine amino acids hints that there may be cryptic ligandable pockets on the KD of Lck, and that covalent modification at these sites could be leveraged to perturb essential structural or regulatory features, providing a mechanistic basis for future allosteric inhibitor development efforts aimed at selectively targeting these amino acids with covalent small molecules. Importantly, deep mutational scanning can reveal sites where covalent liganding is predicted to impact protein function, potentially revealing cryptic ligandable pockets on signaling proteins.

By integrating our activity profiling data with our inhibitor sensitization screens, we identified Lck variants that combine enhanced catalytic output with selective pharmacological control. The I312S Lck variant exemplifies this dual functionality, exhibiting increased phosphotransferase activity and marked sensitivity to nilotinib. When expressed in Jurkat cells, this variant facilitates TCR signaling and, upon drug treatment, functions as an off-switch that suppresses downstream activation even in the presence of endogenous WT Lck. More generally, these results illustrate how multidomain signaling enzymes can be rationally engineered across multiple functional properties: signal amplification and tunable inhibition via clinically approved small molecules. The ability of inhibitor-sensitized Lck variants to perform in cells expressing endogenous WT Lck is particularly notable for translational applications.

Although this study focuses on Lck, we speculate that the principles underlying our approach could be applied across enzyme families and signaling pathways. Many signaling proteins, including other multidomain protein kinases, phosphatases, ubiquitin ligases, and small-GTPases, are regulated by conformational equilibria and long-range allosteric networks that couple enzymatic and regulatory domains. Mutational mapping offers a scalable route to identify function-preserving (or-enhancing) substitutions that simultaneously introduce sensitivity to covalent or reversible ligands. Our strategy enables the selective chemical control of cell signaling outputs without the need for protein fusions with exogenous domains or irreversible genetic modifications. As deep mutational scanning continues to expand in scope, throughput, and model sophistication, we anticipate that similar strategies can be applied to engineer small molecule-gated control over diverse signaling pathways.

## Acknowledgments

This work was supported by the National Institutes of Health (F31GM142170 to Z.E.P., RM1HG010461, UM1HG011969, and R35GM152106 to D.M.F.; R01GM086858 to D.J.M.). Thecontent is solely the responsibility of the authors and does not necessarily represent the official views of the National Institutes of Health.

## Materials and Methods

### Antibodies

| Reagent | Manufacturer | Identifier |
|---|---|---|
| Mouse anti-Lck | Cell Signaling Technologies | Cat# 2657, RRID: AB_21363 |
| Rabbit anti-Phospho-Src Family (Tyr416) | Cell Signaling Technologies | Cat# 6943, RRID: AB_10013641 |
| Mouse anti-ZAP70 | Cell Signaling Technologies | Cat# 2709, RRID: AB_2257531 |
| Rabbit anti-pY319 ZAP70/pY352 Syk | Cell Signaling Technologies | Cat# 2701, RRID: AB_331600 |
| Rabbit anti-pY220 LAT | Cell Signaling Technologies | Cat# 20172, RRID: AB_3073971 |
| Pan phosphor-tyrosine (4G10) | Cell Signaling Technologies | Cat# 96215, RRID: AB_3096295 |
| Monoclonal mouse anti-human CD3 (clone OKT3) | Bio X Cell | Cat# BE0001-2, RRID: AB_1107632 |
| Monoclonal mouse anti-human CD28 (clone 9.3) | Bio X Cell | Cat #BE0248, RRID: AB_2687729 |
| GAPDH | Cell Signaling Technologies | Cat# 2118, RRID: AB_561053 |
| anti-mouse IRDye 800CW | LI-COR | Cat# 926-32210, RRID: AB_621842 |
| anti-rabbit IRDye 680RD | LI-COR | Cat# 926-68071, RRID: AB_10956166 |

### Yeast growth assay

Yeast codon-optimized full-length Lck and indicated mutants were transformed into *S. cerevisiae* ABC16 monster yeast using LiAc transformation (Geitz and Schiestl, 2007) and plated on C-Leu plates. Three colonies for each Lck variant were used to inoculate 1 mL overnight liquid cultures in C-Leu media with 3% glucose. Strains were back diluted into C-Leu media supplemented with 3% raffinose and were allowed to double at least once. Samples were diluted to an OD_600_ of 0.01 in C-Leu media supplemented with 3% galactose, and 150 μL per replicate was plated per well of a 96 well plate. Plates were covered with a Breathe-Easy sealing membrane and incubated in a BioTek Synergy plate reader under constant orbital shaking at 30 °C. OD_600_ was measured every 30 min over 48 hr.

### Yeast western blot protocol

Freshly transformed colonies (∼1 week old) were picked and grown in 1 mL C-Leu 2% glucose overnight at 30 °C with shaking at 250 RPM. Next day, cells were centrifuged at 3,000 g for 5 min in culture tubes, and the media was removed. Cells were resuspended in 5 mL C-Leu 3% raffinose and allowed to double at least once. Cells were pelleted at 3,000 g for 5 min, media was removed, and cells were resuspended in 5 mL C-Leu with 2% galactose. 24 hr later, 100 μL aliquots of cells were centrifuged at 3,000 g for 5 min in microcentrifuge tubes and the media was removed. Cells were stored at-80 °C until further analysis. Cell pellets were thawed on ice and lysed with 200 μL SUMEB Buffer (1% SDS, 8 M urea, 10 mM MOPS, 10 mM EDTA, 0.01% bromophenol blue, 5% BME, 1 mM PMSF, pH 6.8, 1X yeast protease inhibitor (Sigma), 1X phosphatase inhibitor cocktails 2 and 3 (Sigma)). 150 μL glass beads (0.4-0.6 mm) were added to each sample and incubate at ∼1,600 rpm for 10 min at room temperature, then an additional 10 min at 65 °C. Lysate was transferred to a new microcentrifuge tube and centrifuged at 17,000 g at 4 °C for 5 min. Samples were then resolved on SDS-PAGE gels (Criterion, BioRad) and transferred to nitrocellulose membranes (TransblotTurbo, BioRad). Western blots were performed using the indicated primary antibodies and developed using LiCor secondary antibodies.

### Lck Variant Library

Lck single amino acid variant library was commercially synthesized by Twist Biosciences (South San Francisco, CA) and subassembled with random 18 nucleotide barcode into the p415 vector backbone. Synonymous codons were used where possible, and stop codons were included. Following subassembly, the pooled, barcoded variant library was transformed into *E. coli* and serial dilutions were used to estimate library size. Pools of diluted library were collected and analyzed by Illumina sequencing for enumerate library size aiming for an average of 30 barcodes per variant (∼150,000 barcodes total).

### Lck variant library transformation into yeast

Library transformation into yeast was performed following the protocol from Ahler, et al., 2019. Briefly, a single colony of ABC16 monster yeast was picked and grown in 5 mL YPED overnight at 30 °C with shaking. The saturated culture was diluted into 50 mL YPED at an initial OD_600_ of 0.3 and grown until reaching an OD_600_ of ∼2. The culture was centrifuged at 3,000 g for 5 min, the cells were washed with 25 mL sterile H_2_O three times and distributed among 10 microcentrifuge tubes following the last wash. Cells were pelleted at 17,000 g for 30 sec and resuspended in transformation mix containing 240 μL 50% PEG, 50 μL 2 mg/mL salmon sperm (Invitrogen), 36 μL 1 M LiAc. Eight of the samples had 500 ng Lck variant library added, one had 250 ng Lck variant library and 250 ng pRS411 plasmid, and one received only H_2_O without plasmid. All samples were incubated at 42 °C for 45 min after which cells were pelleted and washed with 1 mL sterile H_2_O. The eight library samples were pooled, added to 50 mL C-Leu 2% glucose media and grown with shaking at 30 °C. Serial dilutions of the culture were plated on C-Leu plates to estimate library size. Dilutions of the sample that received the Lck variant library and the pRS411 plasmid were plated on C-Leu-Met plates to estimate the number of double transformants. After 72 hr outgrowth, aliquots of the library were frozen at-80 °C in 20% glycerol.

### Selection and sequencing

The protocol from Ahler, et al., 2019 was followed with modifications made for drug screens. Briefly, aliquots of the frozen Lck variant library were thawed and grown overnight in C-Leu with 2% glucose. The next morning, the cultures were back diluted to OD_600_ 0.5 in C-Leu with 3% raffinose and allowed to double at least once. Selection was initiated by inoculation at OD_600_ 0.01 in 250 mL C-Leu with 2% galactose. Samples of pre-selected cultures were saved and served as timepoint = 0. Cells grew overnight and timepoints were collected at OD_600_, representing approximately 4, 6 and 8 doublings. At each timepoint, 4 OD units of cells were pelleted and stored at-80 °C until further analysis. For drug selections, desired drug concentrations were added when cultures were back diluted to OD_600_ 0.01.

### Lck variant library growth assay analysis

Illumina reads were demultiplexed using bcl2fastq and ea-utils. Activity scores for Lck variants were calculated using Enrich2 (Rubin, et al., 2017). Barcoded Variant SeqLib analysis was used with “weighted least squares” scoring option enabled with “wild type” normalization. Additionally, FASTQ mode parameters included an average quality of 30, and a minimum count of 10. The OD_600_ readings of each timepoint were multiplied by 100 and used as the timepoints. The barcode-variant map from the subassembly was used.

### Cell culture

Jurkat, JCaM1.6, and HEK293T cells were obtained from ATCC and cultured at 37 °C with 5% CO₂. Jurkat and JCaM1.6 cells were grown in RPMI 1640, and HEK293T cells were grown in DMEM, each supplemented with 10% FBS and 2 mM L-glutamine.

### HEK293T drug treatments

500μL HEK293T cells at 0.1×10^6^ cells/mL were plated in a 24 well tissue culture plate. 24 hr later, cells were transfected with Lck variants. Transfection mix was made using 250 ng plasmid DNA, 37 μL serum free DMEM, and 0.75 μL PEI (1 mg/mL). Transfection mixes were incubated at room temperature for 15 min before being added dropwise to cells. 24 hr later, indicated drug at indicated concentration (or DMSO) was added. Cells were incubated in a humidified incubator at 37 °C with 5% CO_2_ for 4 hr. The media was the aspirated, cells were washed once with ice cold PBS, and then cells were lysed directly in 300 μL 1.8X SDS-PAGE loading dye (5X Loading dye: 250 mM Tris, pH 6.8, 45% glycerol, 5% SDS (wt/vol), 0.5 M DTT, 1.86 mM Bromophenol blue). Samples were then collected into microcentrifuge tubes and boiled at 98 °C for 10 min before being resolved via SDS-PAGE (4 μL sample loaded per well) (Criterion gels, BioRad) and transferred to nitrocellulose membranes (TransblotTurbo, BioRad). Westerns were probed with the corresponding primary antibodies, and then developed using LiCor secondary antibodies.

### Lentivirus production

Lentiviral particles were produced in HEK293T cells seeded at 0.3×10⁶ cells per well of a 6-well dish one day before transfection. Cells were co-transfected with the transfer plasmid encoding WT, I312S, or L275R-S323C Lck, the packaging plasmid psPAX2, and the envelope plasmid pMD2.G at a 10:10:1 mass ratio (2.1 µg total) using polyethylenimine (6.3 µg per transfection). The virus-containing supernatant was collected at 48 h post-transfection, cleared by 0.45 µm filtration, and used fresh.

### Transduction

Jurkat cells were seeded at 1×10⁶ cells per well in 6-well plates in virus-containing medium (50%) with and returned to the incubator. The medium was replaced with fresh complete RPMI containing 2 µg/mL puromycin after 48 h. Cell were selected for 7 days, maintained in selection medium, and expanded before use in experiments.

### Categorization of regulatory elements and structural motifs

To classify activity scores and sensitization events by position, we used the following definitions extracted from several sources (Knighton, et al., 1991; Kornev, et al., 2006; Kornev, et al., 2008).

#### Hydrophobic spine positions

408, 409, 424, 432, 433, 428, 324, 370, 371, 372, 259, 271, 422, 362, 383, 276, 288, 273, 292, 303

| Structural motif (number of positions in motif) | Motif start position | Motif end position |
|---|---|---|
| β1 strand (7) | 245 | 251 |
| P-loop (5) | 252 | 256 |
| β2 strand (7) | 257 | 263 |
| β3 strand (9) | 267 | 275 |
| αC-helix (13) | 282 | 294 |
| β4 strand (4) | 304 | 307 |
| β5 strand (4) | 313 | 316 |
| Hinge (7) | 317 | 323 |
| αD-helix (5) | 324 | 328 |
| αE-helix (21) | 338 | 358 |
| β7 & β8 strands (11) | 370 | 380 |
| Activation loop (28) | 381 | 408 |
| αF-helix (17) | 419 | 435 |
| αG-helix (10) | 446 | 455 |
| αH-helix (10) | 468 | 477 |
| αI-helix (15) | 487 | 501 |
| β2 strand-β3 strand loop (3) | 264 | 266 |
| β3 strand-αC-helix loop (6) | 276 | 281 |
| αC-helix-β4 strand loop (9) | 295 | 303 |
| β4 strand-β5 strand loop (5) | 308 | 312 |
| αD-helix-αE-helix loop (9) | 329 | 337 |
| αE-helix-β7&β8 strand loop (11) | 359 | 369 |
| Activation loop-αF-helix loop (10) | 409 | 418 |
| αF-helix-αG-helix loop (10) | 436 | 445 |
| αG-helix-αH-helix loop (12) | 456 | 467 |
| αH-helix-αI-helix loop (9) | 478 | 486 |

### Endogenous Nucleophilic Position Solvent Accessibility Analysis

Solvent accessibility of endogenous nucleophilic amino acids Cys and Lys were assessed using the crystal structure of Lck (PDB ID: 3AD6; Sogabe, et al., 2012). The script FindSurfacePositions was downloaded from the PyMOL Wiki (https://pymolwiki.org/index.php/FindSurfacePositions) and used with its default settings (2.5 Å^2^ cutoff). After positions were identified by the script, they were visually inspected to ensure that the sidechain of each amino acid were solvent exposed.

### Amino acid position distance from ATP-binding site

To calculate the distance of each amino acid’s α-carbon from the ATP-binding site, we first recorded the coordinates of the center of mass (COM) of imatinib bound to the active site of Lck (PDB ID: 2PL0; Jacobs, et al., 2008). We then used PyMol’s distance function to calculate the distance between the COM and each amino acid’s α-carbon.

### Principal component analysis

Principal component analysis (PCA) was performed to reduce the dimensionality of the inhibitor sensitization dataset and identify the dominant sources of variance among sensitization events. Prior to analysis, we removed variants that were classified as deactivating in our activity screen, and variants that were not quantified in all five inhibitor datasets yielding a total of 2,093 variants for analysis. PCA was performed in GraphPad Prism and the number of principal components retained was determined by parallel analysis (default settings: 1000 simulated datasets; 95th percentile cutoff). Samples were projected onto the first two principal components (PC1 and PC2) for visualization, and the percentage of variance explained by each component is reported on the corresponding axes. Feature loadings were examined to identify the variables contributing most to each component.

### Relative Solvent Accessible Surface Area (rSASA) calculations

Accessible surface areas were downloaded from the Genomics 2 Proteins (G2P) portal (Kwon, et al., 2024), calculates surface area from AlphaFold models. Absolute accessible surface areas were normalized to standard amino acid surface areas, summarized in the table below, to yield the relative Solvent Accessible Surface Area (rSASA) for each residue. These rSASA values were used to calculate the percentage of Lck’s solvent-exposed surface area covered by WT-like Cys and Lys substitutions.

| Residue: Solvent Accessible Surface Area |
|---|
| Ala: 115 |
| Arg: 225 |
| Asn: 160 |
| Asp: 150 |
| Cys: 135 |
| Gln: 180 |
| Glu: 190 |
| Gly: 75 |
| His: 195 |
| Ile: 175 |
| Leu: 170 |
| Lys: 200 |
| Met: 185 |
| Phe: 210 |
| Pro: 145 |
| Ser: 115 |
| Thr: 140 |
| Trp: 255 |
| Tyr: 230 |
| Val: 155 |

### Steric Perturbation Index Calculations

Identifying positions intolerant of substitution reveals amino acids important for catalysis and/or folding, where covalent modification would likely inactivate Lck, but it does not directly predict the functional consequence of a ligand occupying an adjacent binding pocket. We developed the Steric Perturbation Index (SPI) to model this scenario, reasoning that bulky aromatic substitutions (Phe, Trp, and Tyr) best approximate the steric footprint of a covalent ligand engaging a pocket adjacent to a nucleophilic amino acid, while more modest substitutions (Ala, Thr, Ser, and Cys) report on the functional consequences of losing the WT amino acid side chain itself. Positional SPI values (PSPI) were calculated as the difference between the average activity scores of these two substitution classes at each solvent-exposed Lys position, with a negative or positive PSPI predicting that covalent engagement at that site would inhibit or activate kinase activity, respectively. We also modeled the scenario where bulky substitutions to residues surrounding an endogenous Lys are inhibitory, termed Surrounding SPI scores (SSPI). These scores were derived by calculating the SPI scores for every position within 5 Å of each solvent-exposed endogenous Lys, taking the average of these scores as the SSPI score. Negative SSPI scores suggest that liganding at these sites could result in phosphotransferase inhibition.

### Drug Sensitization Scores

The Lck variant library was transformed into yeast, and a sample was collected prior to Lck induction. Lck expression was then induced in the presence of either DMSO or each drug at the concentration that gave minimal but detectable inhibition of WT Lck (12.5 µM imatinib, 250 nM nilotinib, 2 µM dabrafenib, 17 µM erlotinib, 8 µM gefitinib), and samples were collected throughout outgrowth. Plasmids were extracted, and activity scores for each inhibitor were calculated from two independent biological replicates as described above for the activity profiling experiments. For each variant, we then calculated a “sensitization score” as its drug-treated activity score minus its DMSO activity score, such that more negative scores indicate greater drug sensitization. Drug sensitized variants, or “sensitization events,” were defined as those variants with sensitization scores more than 2 standard deviations below the mean sensitization score for each drug.

### TCR Activation Immunoblot Analyses in Jurkat and JCaM1.6 Cells

Parental Jurkat or JCaM1.6 cells or those recombinantly expressing WT, I312S, or L275R-S323C Lck were normalized to 1×10^6 cells/mL in serum free RPMI media and pre-treated with or without nilotinib (1 µM, 1 h). Cells were then stimulated with anti-CD3:CD28 antibodies (5:2 µg/mL, 1 min), quenched on ice, lysed with RIPA buffer containing protease and phosphatase inhibitors, and subjected to SDS-PAGE and immunoblot analysis.

### IL-2 ELISA

Jurkat cells, or Jurkat cells recombinantly expressing WT or I312S were pre-treated with DMSO or nilotinib (1 µM, 1 h), followed by stimulation with anti-CD3:CD28 antibody cocktail (5:2 µg/mL, 6 h). After quenching on ice, IL-2 levels in the supernatant were quantified using the Human IL-2 ProQuantum Immunoassay Kit (Catalog # A35603) according to the manufacturer’s protocol.

### Hierarchical clustering of positions with 5 or more activating substitutions

Atomic coordinates of the α-carbons from positions harboring 5 or more activating mutations were extracted from the AlphaFold model of Lck (AlphaFold: AF-P06239-F1) and clustered using the hclust() R package.

### Structural alignment of Lck and EGFR

The AlphaFold model of Lck (AlphaFold: AF-P06239-F1) was aligned to the crystal structure of neratinib bound to EGFR T790M/L858R (PDB: 3W2Q; Sogabe, et al., 2012) using Pymol (version 1.3).

## Author Contributions

Z.E.P., D.M.F., and D.J.M. conceived the study. Z.E.P. developed the methodology, performed the activity, inhibitor sensitization and validation experiments, analyzed the data, and generated the figures. M.A.C. helped design the barcoded Lck library. C.T.W. performed Illumina sequencing. A.N.G.. J.L.B., and J.C. performed yeast validation experiments. A.F.A. and J.J.S. planned and performed the experiments and generated and validated the cell lines. J.J.S., N.A.P., D.E.U., and D.S.B. provided experimental support. Z.E.P. and D.J.M. wrote the original manuscript draft.

---

**Figure S1.**
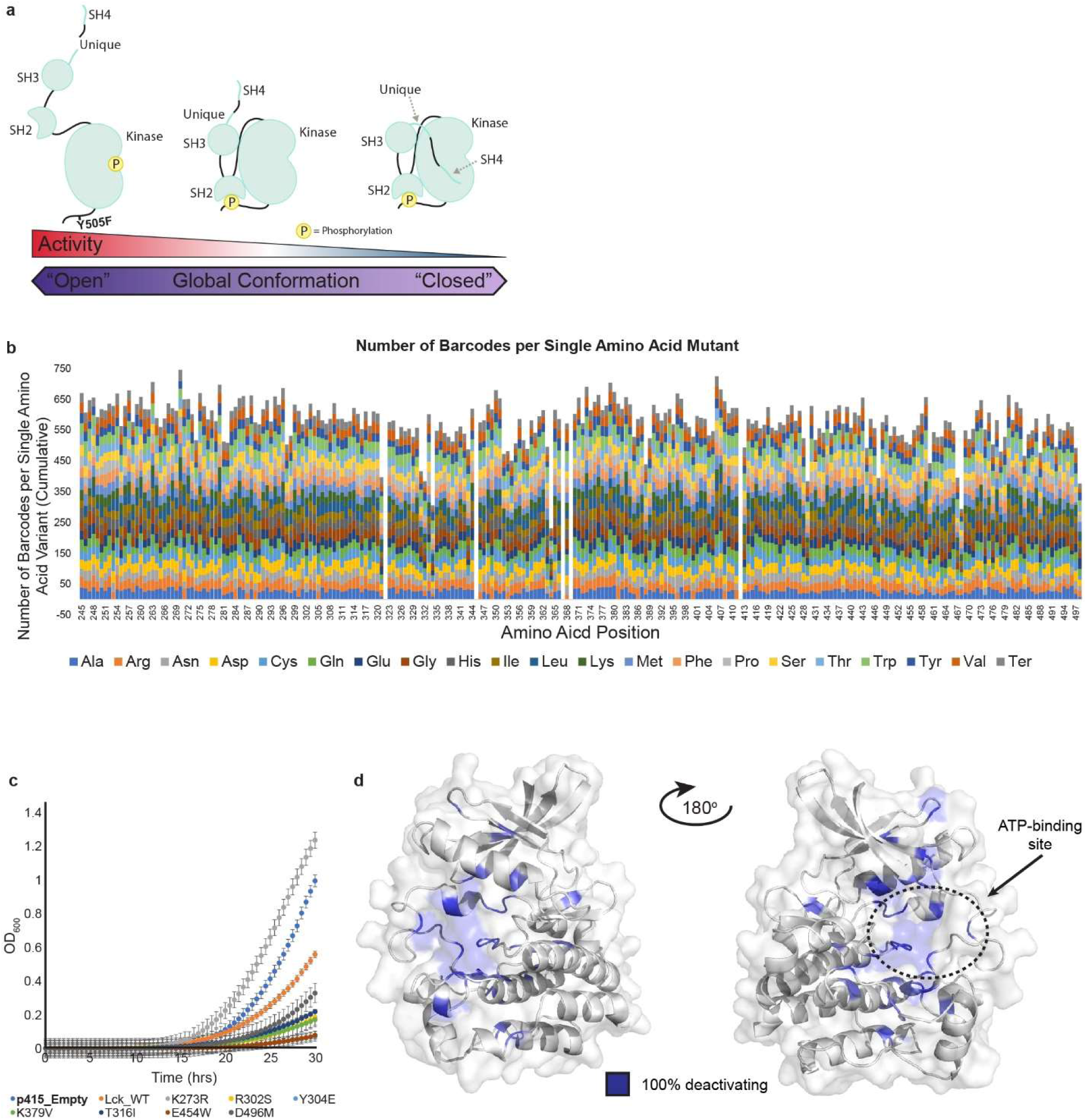
Profiling Lck’s Phosphotransferase Activity in Yeast. (A) Conformational equilibrium of Src Family Kinases and corresponding phosphotransferase activities. (B) Summary of the PacBio sequencing data of the Lck variant library. Stacked bars represent the number of DNA barcodes at each position associated with each amino acid substitution at that position. (C) Individually assessed growth curves for yeast expressing Lck WT, K273R (kinase dead), R302S,Y304E, K379C, T316I, E454W, D496M, or a control vector. Datapoints represent average ± s.e.m. of three biological replicates. (D) Amino acid positions where 100% of variants are classified as deactivating mapped onto the AlphaFold model of Lck’s KD (AlphaFold: AF-P06239-F1).

**Figure S2.**
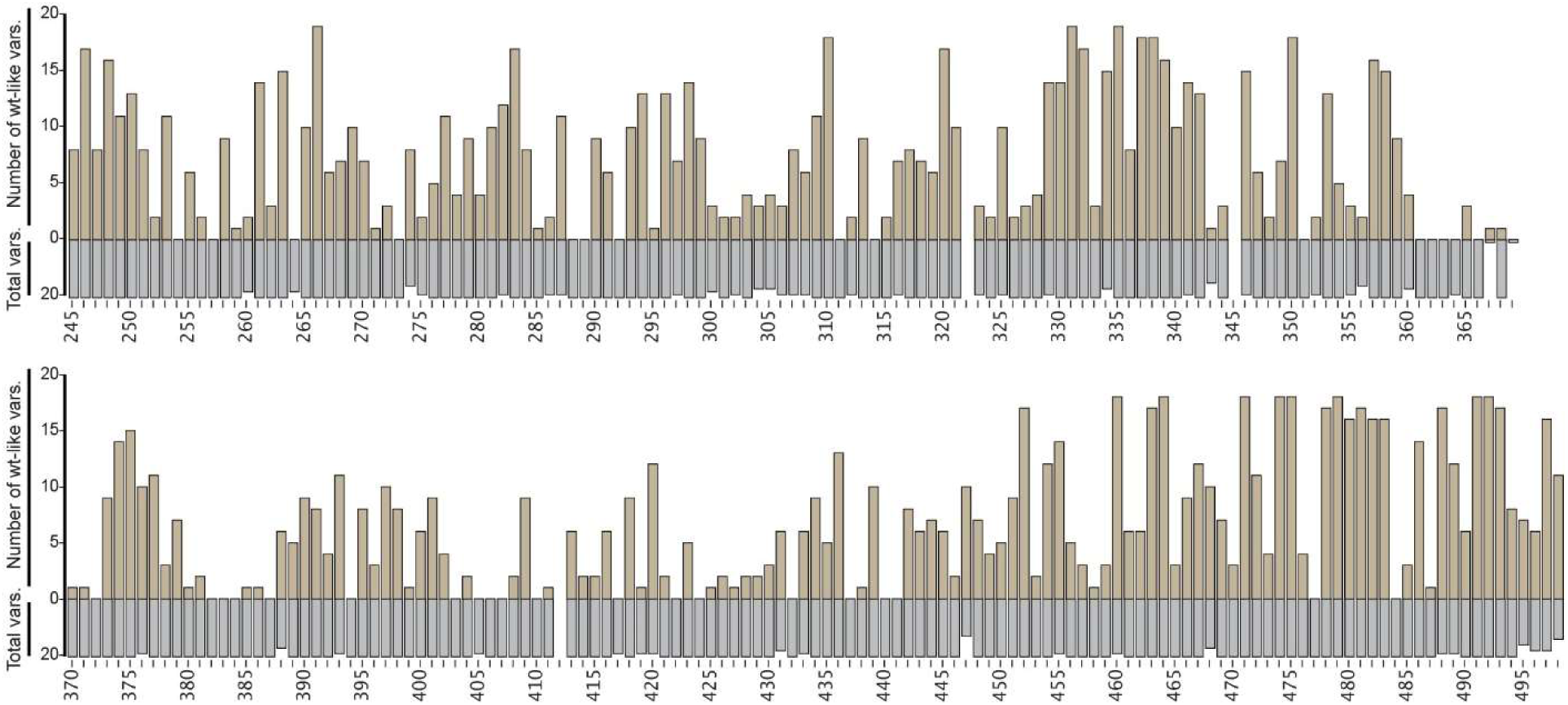
WT-like amino acid substitutions on Lck’s KD. (A) Number of WT-like amino acid substitutions per position. Tan bars (upper) represent the number of WT-like variants at a given position on Lck’s KD. Grey bars (lower) represent the total number of variants quantified at that position.

**Figure S3.**
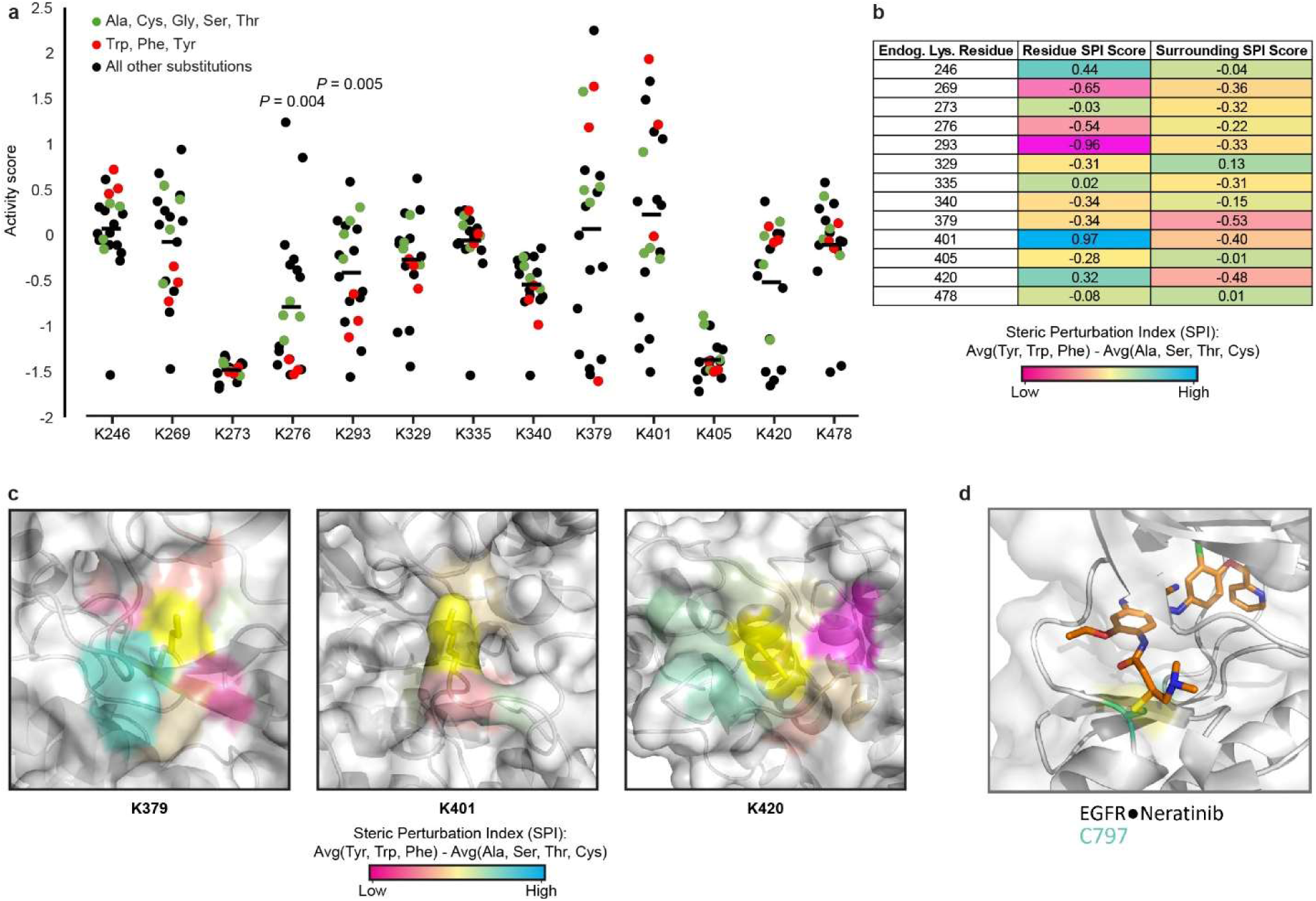
Activity scores of endogenous lysine positions and the covalent adduct between EGFR and neratinib. (A) Activity scores for all solvent accessible endogenous lysine variants. (B) Positional and Surrounding Steric Perturbation Index scores for all endogenous lysine amino acids. SPI scores are calculated as the average of Tyr, Phe, and Trp activity scores minus the average of Ala, Ser, Thr, Cys activity scores. Positional SPI scores reflect the SPI score for each endogenous lysine amino acid. Surrounding SPI scores reflect the average SPI score for all positions within 5 Å of each endogenous lysine position. (C) Endogenous lysine positions with the most negative Surrounding SPI scores (K379, K401, and K420). Indicated lysine amino acid is shown with side chain in yellow. SPI scores for all positions within 5 Å are shown, shaded according to the color scale. (D) Covalent adduct between the ATP-competitive inhibitor neratinib bound to EGFR T790M/L858R at C797 (PDB: 3W2Q; Sogabe, et al., 2012).

**Figure S4.**
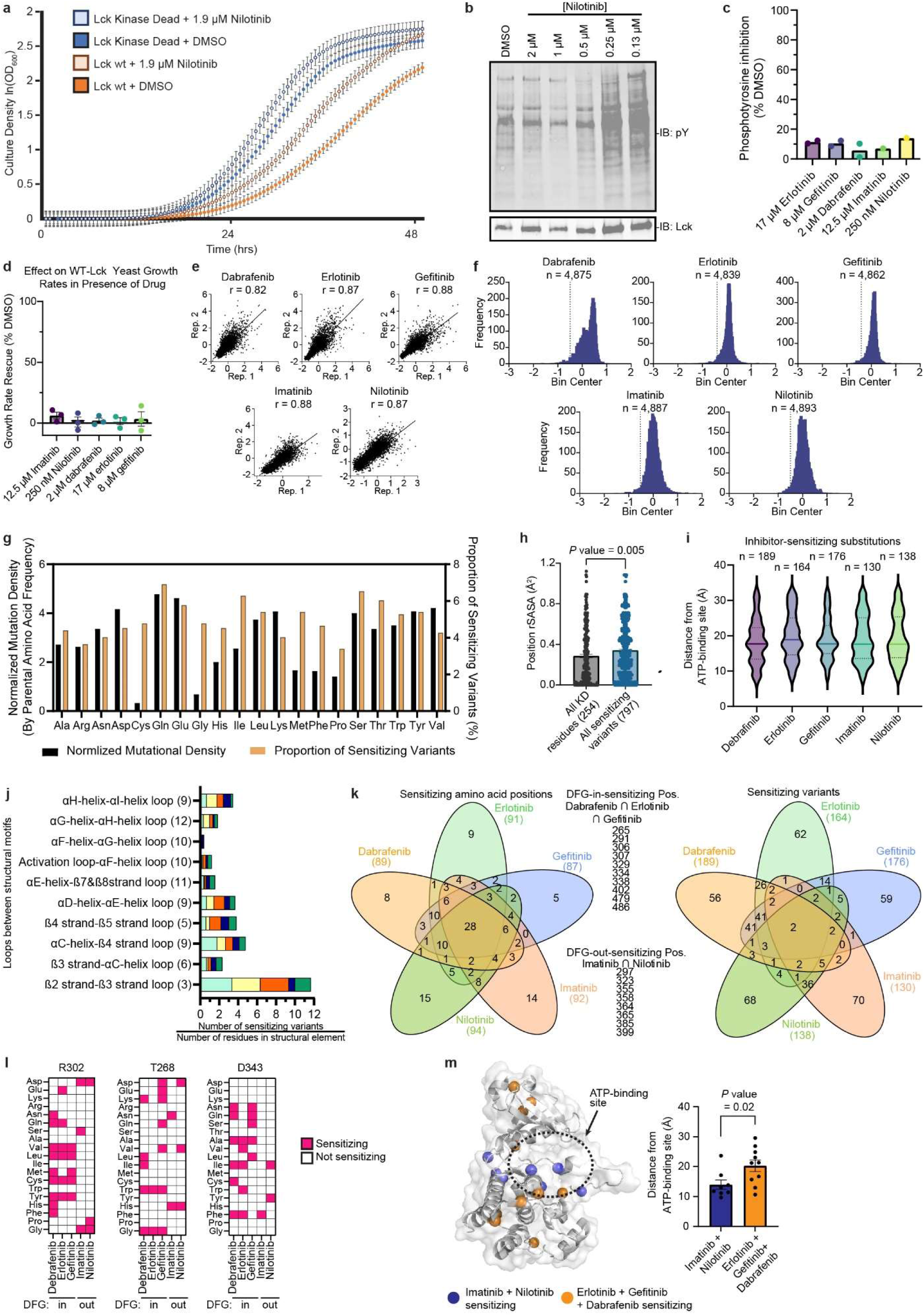
Identification of Drug-Sensitizing Lck Variants using yeast growth-based sensitization assay. (A) Individually assessed growth curves for yeast expressing Lck WT, K273R (kinase dead) grown in the presence of 1.9 µM nilotinib or DMSO. Datapoints represent average ± s.e.m. of three biological replicates. (B) Phosphotyrosine immunoblot (IB) of yeast expressing Lck WT grown in the presence of DMSO or varying concentrations nilotinib. Representative of experiments performed for erlotinib, gefitinib, neratinib, dabrafenib, and imatinib. (C) Quantification of phosphotyrosine blots of yeast treated with indicated inhibitors at indicated concentrations. Phosphotyrosine levels are normalized to Lck levels. These concentrations of each inhibitor were used in bulk yeast growth assay. Datapoints represent two independent biological experiments except for imatinib and nilotinib which are from a single experiment. Bars represent averages. (D) Growth rate rescue of WT Lck-expressing yeast treated with the indicated inhibitor at the indicated concentration. Growth rate rescue calculated as a percentage of growth rate in the presence of DMSO compared to inhibitor treatment. Dots represent individual biological replicates, bars represent averages, error bars represent s.e.m.. (E) Correlation of Lck activity scores between replicates of each drug treatment. (F) Histogram of all sensitivity scores. Vertical dotted line represents the average sensitization score minus 2 standard deviations. Drug sensitized variants (or “sensitizing events”) are those to the left of this dotted line. (G) Amino acid-level analysis of all sensitization events. Normalized mutational density: Counts of all variants for each endogenous amino acid divided by the endogenous frequency of each amino acid in the KD of Lck. Proportion of sensitizing variants: Proportion of all sensitizing events for each amino acid. (H) Bar chart representing average solvent accessibility scores (rSASA) for all sensitizing events compared to all KD positions. Statistical significance assessed using students t test. Bars represent averages ± s.e.m. (I) Distance between α-carbons for all sensitizing events and the center of mass of imatinib bound to Lck (PDB 2PL0; Jacobs, et al., 2008). Solid lines represent median (dabrafenib, 17.7 Å; erlotinib, 18.8 Å; gefitinib, 17.7 Å; imatinib, 17.7 Å; nilotinib, 17.7 Å), dotted lines represent quartiles. The median distance for all 797 sensitizing variants is 17.7 Å from the center of mass of imatinib, with 25% of variants located more than 24.2 Å away. (J) Sensitizing events stratified by the loops between structural motifs of Lck. Bar represents all sensitizing events on a given structural motif divided by the number of positions in that motif. The shaded region of each bar represents the proportion of the total number of sensitizing events on that motif belonging to each inhibitor. (K) Venn diagram representing all inhibitor sensitizing amino acid positions (left) sensitizing events (right) that are shared between inhibitors or unique to each inhibitor. (L) Heatmap showing the R302, T268, and D343 sensitizing events. (M) Analysis of positions with inhibitor class-specific sensitizing events: DFG-in (Erlotinib, gefitinib, dabrafenib) versus DFG-out (Imatinib, nilotinib) inhibitors. Bar chart represents the average distance of sensitizing positions between each inhibitor class from imatinib’s center of mass (PDB 2PL0; Jacobs, et al., 2008). Dots represent individual positions, bars represent averages, error bars represent s.e.m.. Statistical significance determined using Students t test.

**Figure S5.**
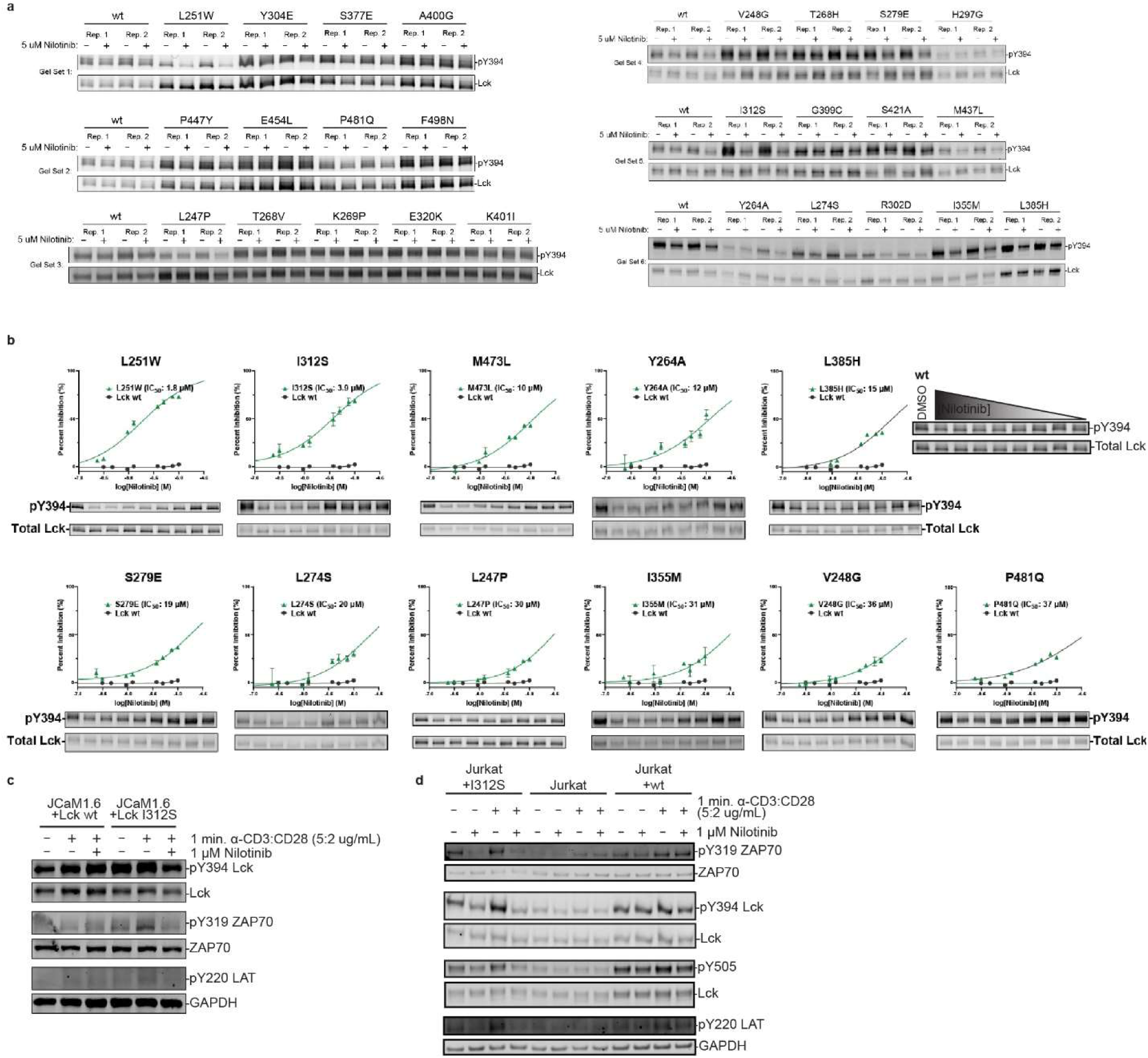
Characterization of Drug-Sensitizing Lck Variants. (A) Immunoblots (IB) for a panel of nilotinib-sensitized variants individually expressed in HEK293T cells treated with 5 μM nilotinib or DMSO for 4 h (n=2). (B) Dose-dependent analysis of nilotinib sensitivity for individual nilotinib-sensitized variants that showed increased levels of inhibition relative to WT Lck when treated with 5 µM nilotinib for 4 h (“performers”). Blots are representative of two replicates. Data points represent averages ± s.e.m. (C) Second replicate of WT and I312S Lck characterization in J.CaM1.6 cells. Immunoblot analysis of JCaM1.6 cells recombinantly expressing WT or I312S Lck that were treated with DMSO or 1 µM nilotinib for 1 h. Cells were then stimulated with anti-CD3:CD28 antibody cocktail for 1 min and then quenched on ice. (D) Second replicate of immunoblot analysis of parental Jurkat cells or those recombinantly expressing WT or I312S Lck pre-treated with or without nilotinib (1 µM, 1 h). Cells were then stimulated with anti-CD3:CD28 antibodies (5:2 µg/mL, 1 min), quenched on ice, lysed and subjected to SDS-PAGE and immunoblot analysis.

